# Pipeline for RNA sequencing data analysis by combination of Nextflow and R

**DOI:** 10.1101/2023.10.15.562329

**Authors:** Jia-Hua Qu

## Abstract

With the development of high-throughput technologies, RNA sequencing (RNA-seq) has become a widely used technology in biological studies and thus a large number of RNA-seq data are emerging and remain to be analyzed. Although there are many different options for analysis methods and tools, a unified pipeline for RNA-seq data analysis is always necessary for a laboratory. Given the update of new methods and tools, I summarized my customized analysis codes to generate an updated pipeline for RNA-seq data analysis. During aging, gene mutations accumulate, and hormone regulation is disrupted, which may exacerbate age-related diseases. Therefore, we generated a dataset from mice with a gene mutation or not and under different hormone treatments to study the effects of two factors, i.e., hormone and gene mutation, on the transcriptome. Based on the Nextflow nf-core rnaseq pipeline, this project established this pipeline consisting of three stages: (1) upstream analysis containing quality control of fastq files before and after trimming, trimming, alignment, and quantification; (2) midstream analysis containing count normalization, differentially expressed genes analysis, and visualization via boxplot, PCA, t-SNE, sample distance heatmap, MA plot, volcano plot, and gene expression heatmap; and (3) downstream analysis containing functional enrichments of KEGG pathways and GO terms. Results showed distinct effects of the single factor as well as interactive effects of the two factors. Codes are also provided for readers who want to customize their analysis pipeline adapted from this pipeline easily.

## Introduction

In the field of modern biological studies, omics are widely used, and thus high-throughput data are generated every day in need of customized analysis. There are many different options for analysis methods and tools. They can be classified into two major types: graphical user interface and command line interface. The former can be divided into website applications and local computer applications; the latter can be divided into programs written in different programming languages, such as Bash, R, Python, Perl, Groovy, etc.

The graphical user interface is convenient for a broad range of scientists who want to make full use of omics but don’t have programming skills to analyze high-throughput data, whereas the command line interface is more flexible in customizing analysis and helps save time in repeated procedures. Website applications are usually updated more frequently than local computer applications, so website applications often provide access to the latest version of databases, but they may be unstable. In contrast, local computer applications are more stable and can be used to reproduce results better than website applications. Partek Flow is a representative website application with a graphical user interface, and it provides convenience in omics data analysis, particularly useful in single-cell RNA-seq data analysis[1]. JMP and GraphPad Prism are two well-used local computer applications with graphical user interfaces with statistical functions as well[2].

Despite the advantages and convenience of the graphical user interface, professional bioinformaticians prefer the command line interface because of its more powerful functions that can also be extendible and scalable. For example, fastq files of a larger number of samples are usually analyzed in the Linux system, because the Linux system can be used to process high-throughput data in parallel and fast[3]. R and Python are more and more popular because both are open-sourced and free[4]. A lot of R packages and Python libraries are available and provide conveniences for bioinformatics analysis.

In addition, there are some tools invented especially for building pipelines, such as Nextflow[5], Snakemake[6], WDL and Cromwell[7], etc. For example, Nextflow is well-used, because it enables scalable and reproducible scientific workflows and allows the adaptation of pipelines written in the most common scripting languages. In addition, Nextflow has a very active user community which released many curated nf-core pipelines (https://nf-co.re/)[8] that can be modified for customized use.

Given a typical pipeline for RNA sequencing (RNA-seq) data analysis consists of multiple steps, this project divided it into three stages: (1) upstream data process; (2) midstream visualization of count matrix; and (3) downstream functional analysis. Different tools have different advantages at different stages and different sets of data require customized analysis, so the combination of different tools is a better choice in a practical project.

Because of the advantages of different tools, the combination of various tools allows us to customize our bioinformatics analysis flexibly[9]. My pipeline was built by the combination of Nextflow nf-core rnaseq pipeline and R with many R packages.

Aging is a heavy burden to the modern society. The recent review proposed ten aspects of biomarkers of aging, including epigenetic changes, genetic instability, telomere shortening, nuclear body disorders, cell cycle arrest, mitochondrial malfunction, proteostatic stress, metabolic alterations, signaling pathway rerouting, and senescence-associated secretory phenotype (SASP)[10]. In the aspect of genetic instability, mutations that are caused by genomic instability and impaired DNA repair exhibit an age-related increase in a tissue-specific manner[11, 12]. In addition, the endocrine and exocrine in multiple organs are closely related to aging[13]. For example, during aging, gene mutations are accumulated, and hormone regulation is disrupted, which may exacerbate age-related diseases in return. I applied this pipeline to our in-house dataset that was generated from mice with a gene mutation or not and under different hormone treatments. My results showed distinct effects of a single factor as well as interactive effects of the two factors, i.e., hormone and gene mutation, on mouse transcriptome.

## Tools

### Linux system

Wynton HPC (https://wynton.ucsf.edu/hpc/index.html) - UCSF Research Computing at Scale, based upon the CentOS 7.

### Pipeline tool

The nf-core/rnaseq pipeline (https://nf-co.re/rnaseq/3.10.1)[8], based upon Nextflow (https://www.nextflow.io/)[5].

### R

R version: 4.1.3 or 4.2.2

RStudio version: 2023.09.0 Build 463

R packages versions:

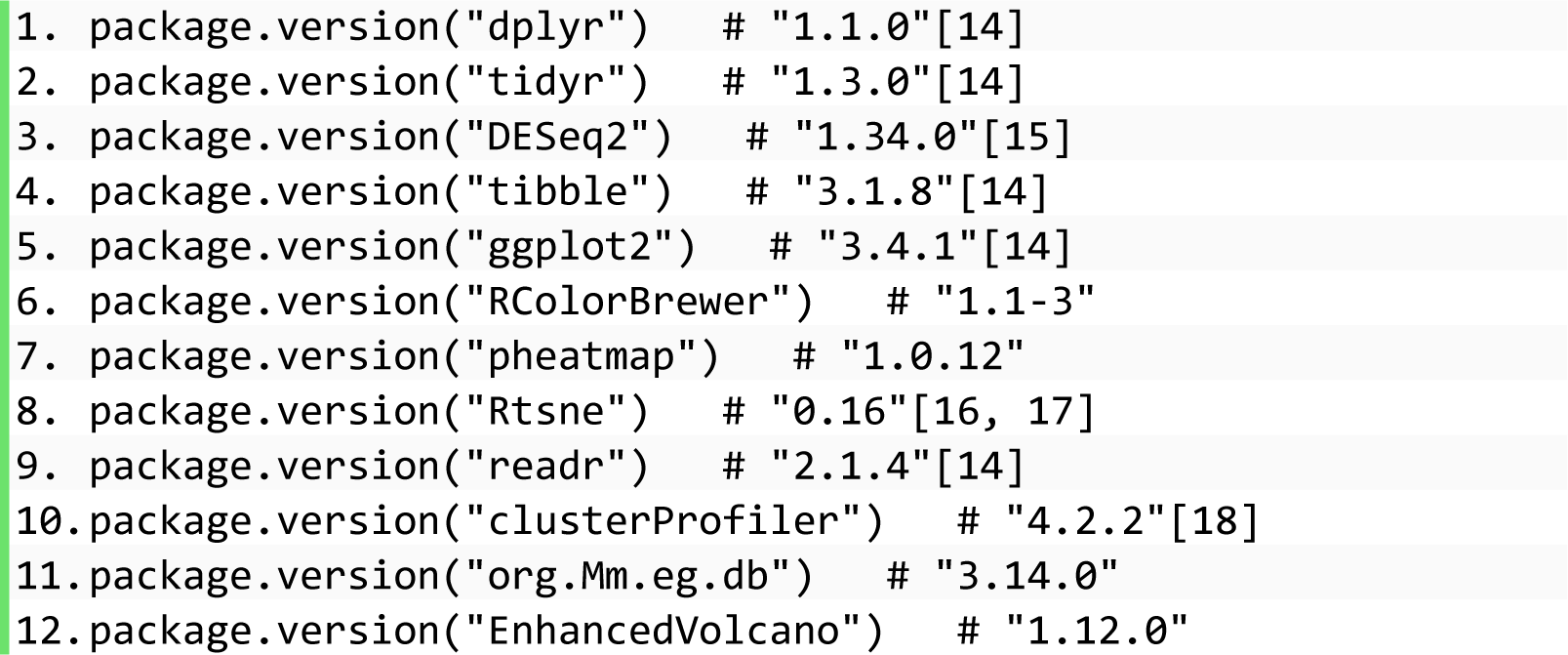

### Dataset

In this project, our in-house RNA-seq dataset was used as a demo. Sequencing type: SE. Library Kit: Nugen Universal Plus. Machine: HiSeq4000. Because our in-house RNA-seq dataset is unpublished and the application of this pipeline is not restricted to this dataset, the detailed meta information is hidden. In brief, this dataset was obtained from mice with a gene mutation (mut) and their wild-type littermates (con), both of whom received treatment of two hormones singly (E2, P4) or combinedly (E2P4) or placebo (Plac). In total, this dataset contains eight groups of samples: Plac_con, Plac_mut, E2_con, E2_mut, P4_con, P4_mut, E2P4_con, and E2P4_mut. This project did not intend to describe the gene mutation within this dataset or to explain the definition of different hormone treatments explicitly. When this pipeline is applied in other projects, *mut* can represent any other gene mutation, and *E2* or *P4* can represent any other internal or external stimuli or disease states.

Based on the Nextflow nf-core rnaseq pipeline, this project established this pipeline (Figure 1) consisting of three stages: (1) upstream analysis containing quality control of fastq files before and after trimming, trimming, alignment, and quantification; (2) midstream analysis containing count normalization, boxplot, PCA, t-SNE, sample distance heatmap, differentially expressed genes analysis, MA plot, volcano plot, and gene expression heatmap; and (3) downstream analysis containing functional enrichments of KEGG pathways and GO terms.

**Figure 1.**
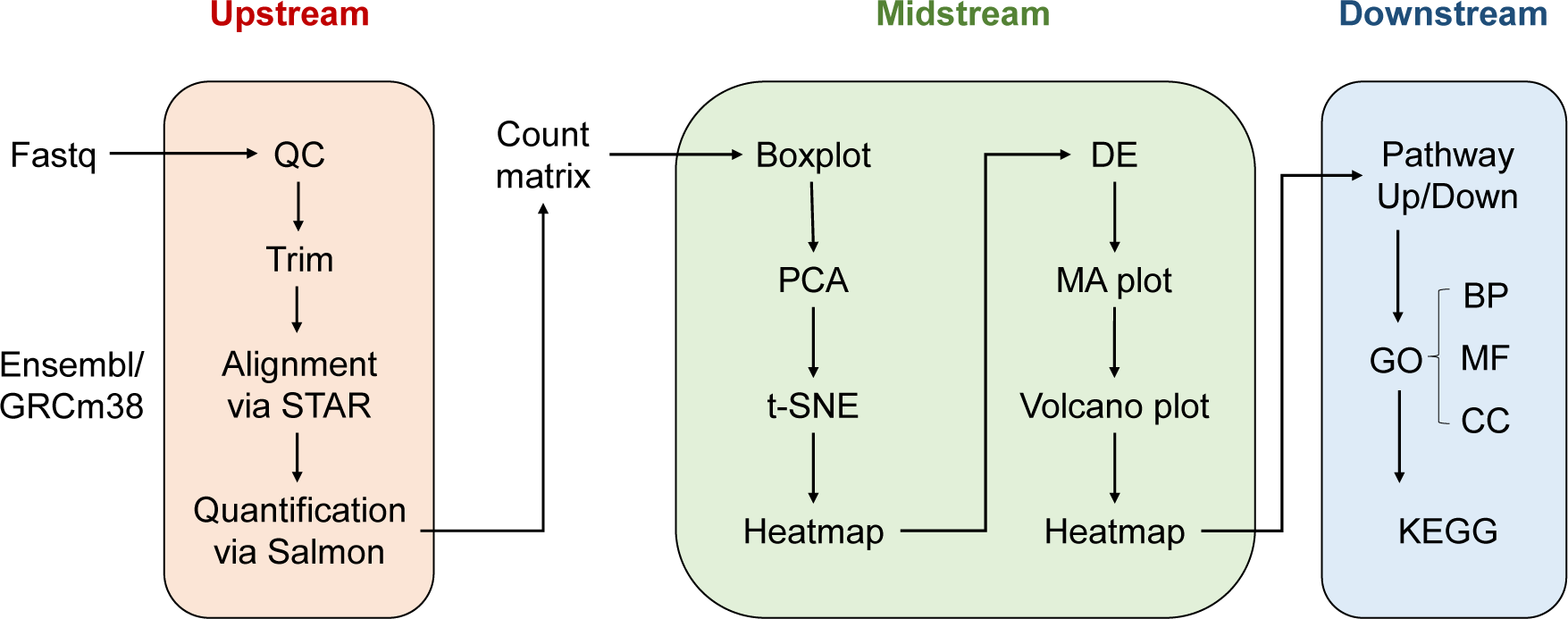
Workflow of RNA-seq data analysis. Upstream analysis of fastq files via Nextflow, midstream analysis of count matrix and differentially expressed genes in R, and downstream analysis of functional enrichment.

### Upstream analysis with Bash and Nextflow

1. Create a conda environment. The virtual environment technique is an ideal way to manage systems and software, so I recommended creating a virtual environment via conda (https://docs.conda.io/en/latest/).

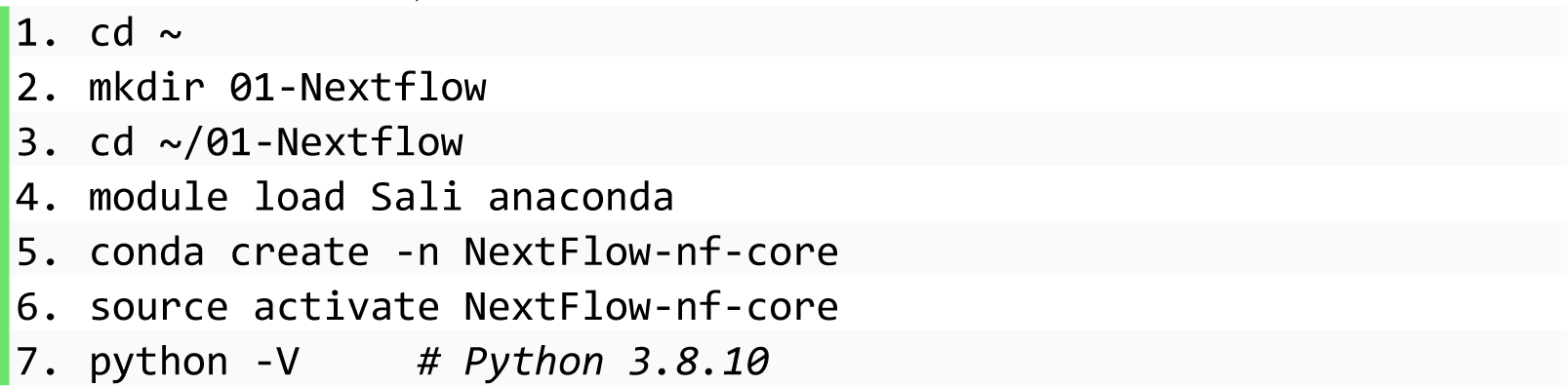
2. Install Nextflow, nf-core, and rnaseq pipeline. Singularity is a scientific container for the mobility of computing[19], so it is strongly recommended to use a singular image of the nf-core/rnaseq program to reproduce the analysis.

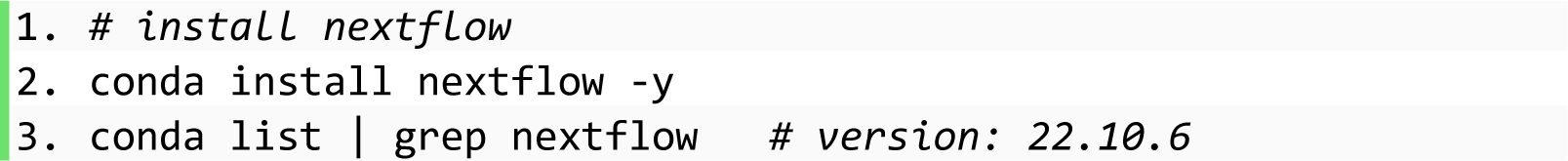
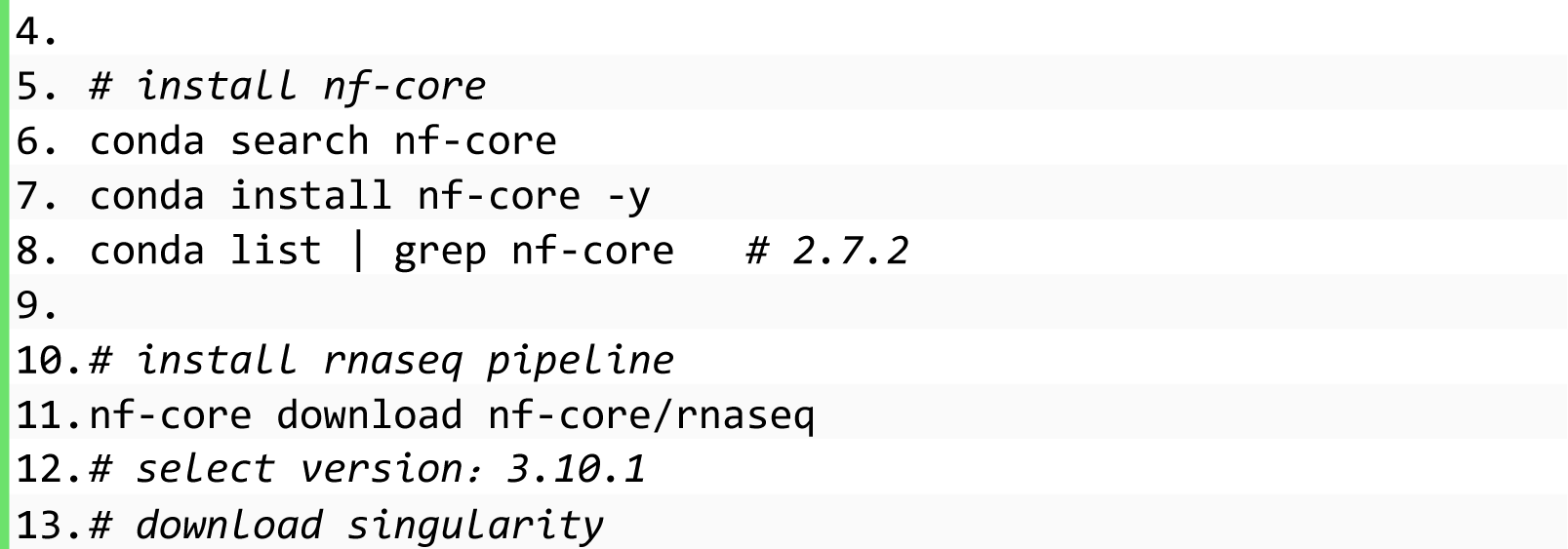
3. Prepare the reference genome. Because the data were obtained from mice, the GRCm38 reference genome and its accompanied annotation were downloaded from iGenome (https://support.illumina.com/sequencing/sequencing_software/igenome.html) and used in this project.

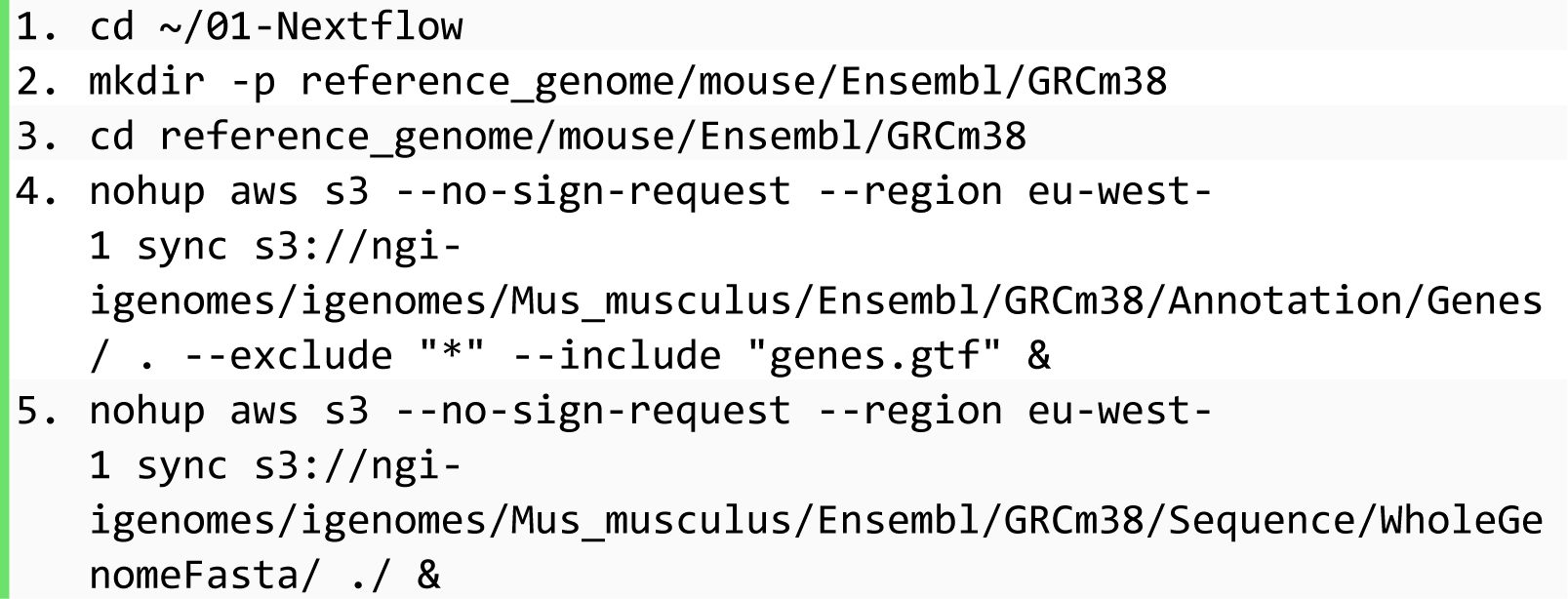
4. Prepare fastq files.

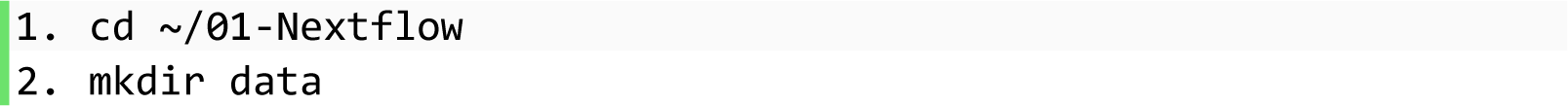 Copy fastq files that need to be analyzed into the folder ∼/01-Nextflow/data.
5. Make a sample sheet. Download the Python script to make a sample sheet for the subsequent analysis.

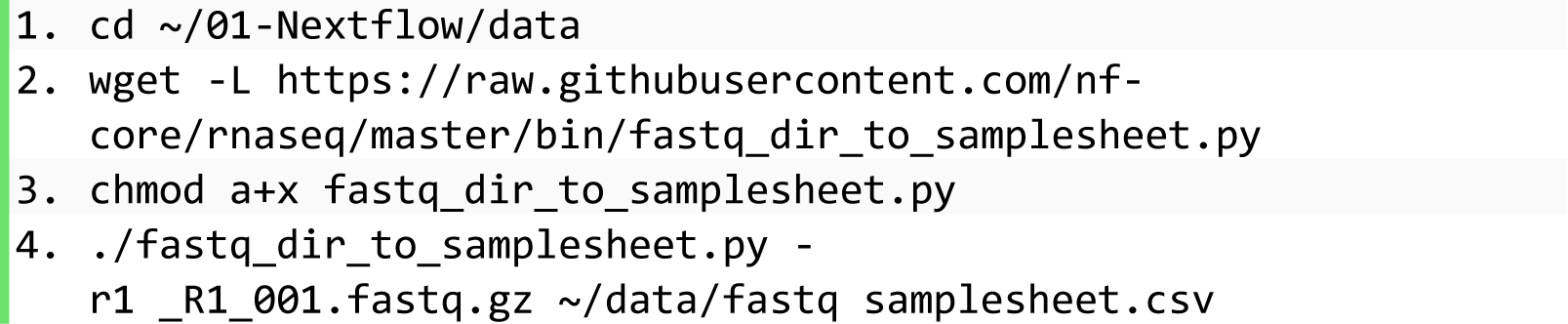
6. Customize a config file.

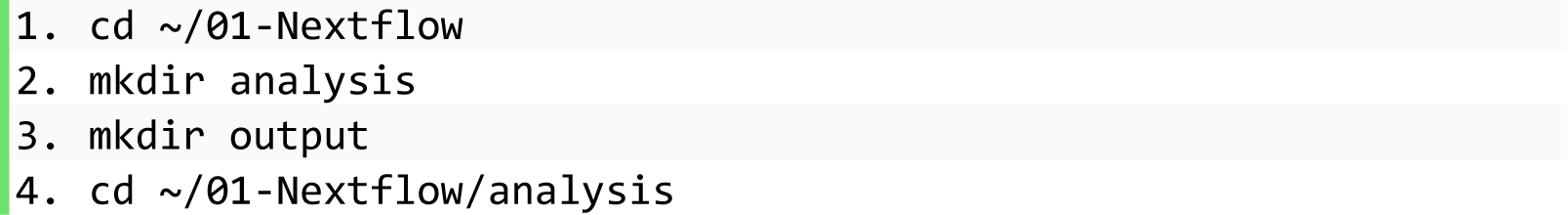

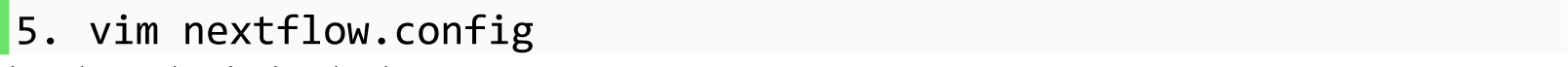 Customized analysis includes:

- FastQC – Raw read QC
- TrimGalore – Adapter and quality trimming
- STAR – Alignment
- Salmon – Quantification
- RSeQC – Various RNA-seq QC metrics
- DESeq2 – PCA plot and sample pairwise distance heatmap and dendrogram
- MultiQC – Present QC for raw reads, alignment, read counting, and sample similarity In addition, in the config file, we specified the paths to the input sample sheet of fastq files to be analyzed, reference genome files, and folder to save the output.

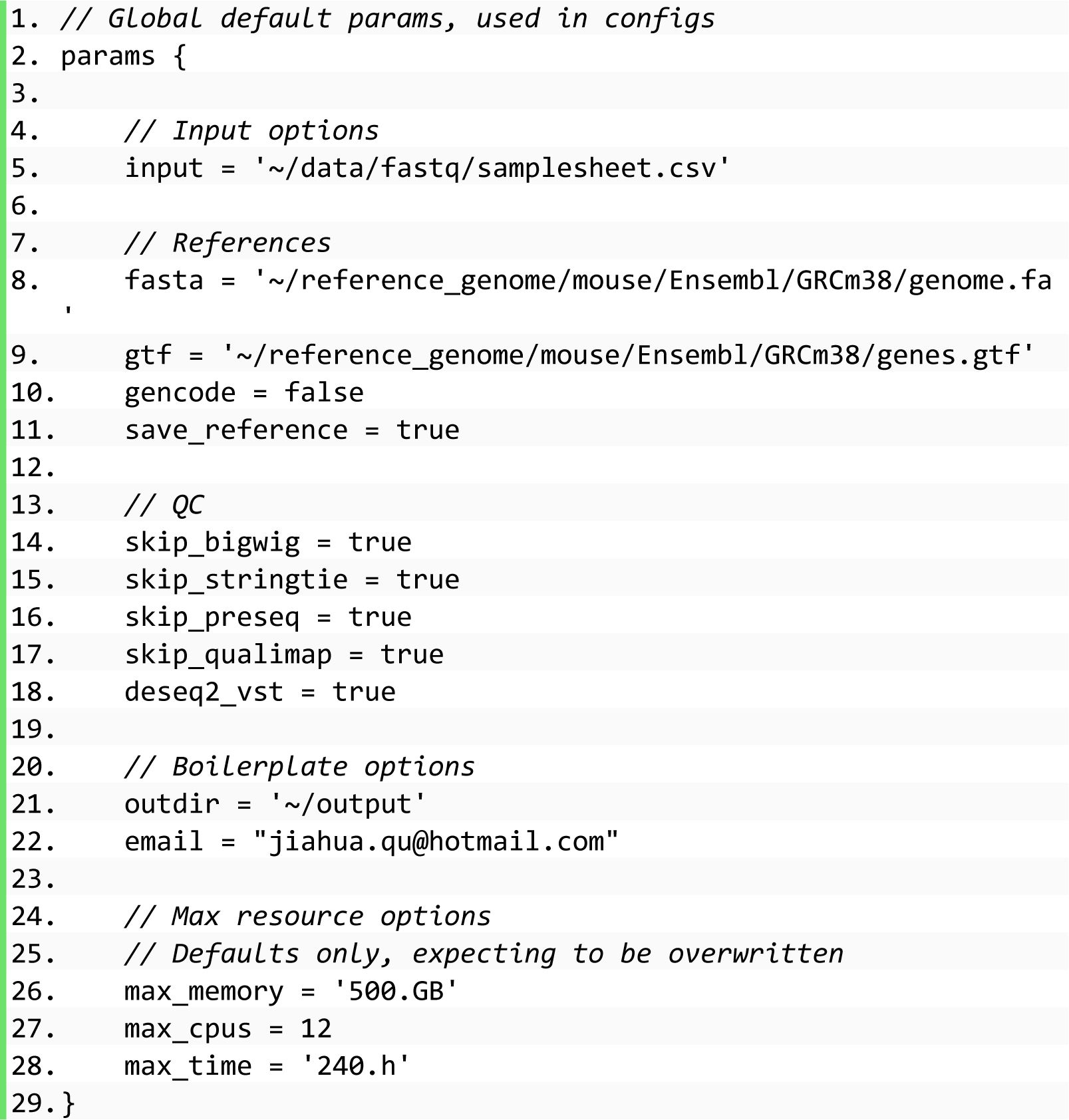
7. Make a bash script.

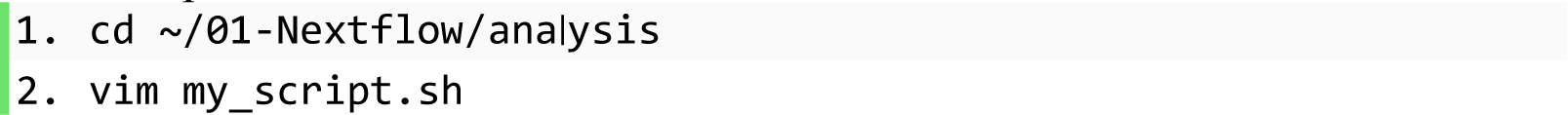 In the bash script, we specified: (1) required computation resources including core number, memory, and running time; (2) paths to input files including log file that records progress, Nextflow workflow program, singularity profile, and config file with customized settings.

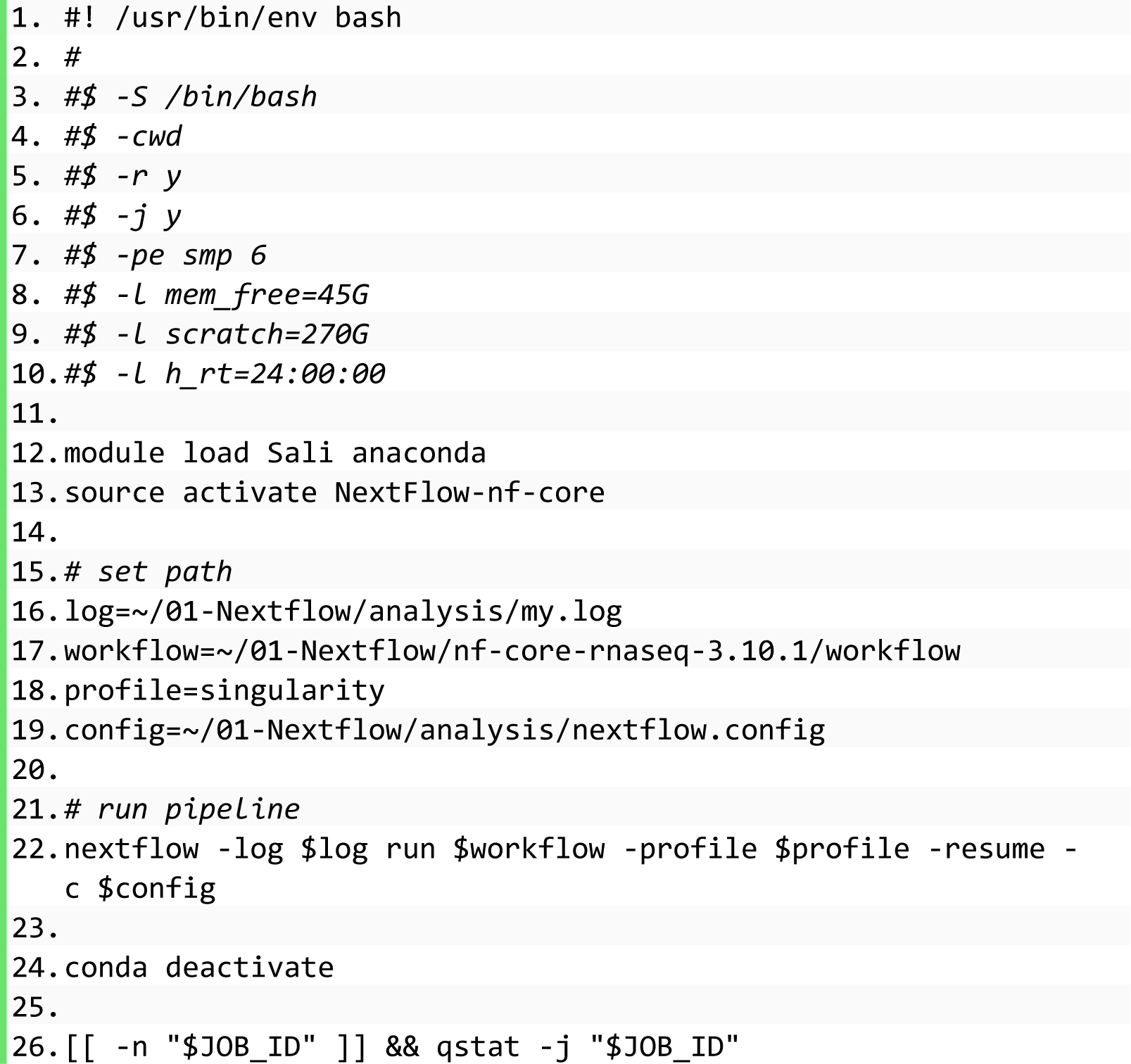
8. Submit a job to run in the background.

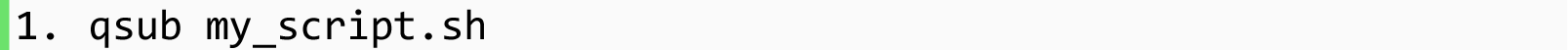
9. Output. The output contains two major parts: (1) star_salmon output including count matrix and metadata, and (2) MultiQC report showing quality control of fastq files before and after trimming (Figure 2).

**Figure 2.**
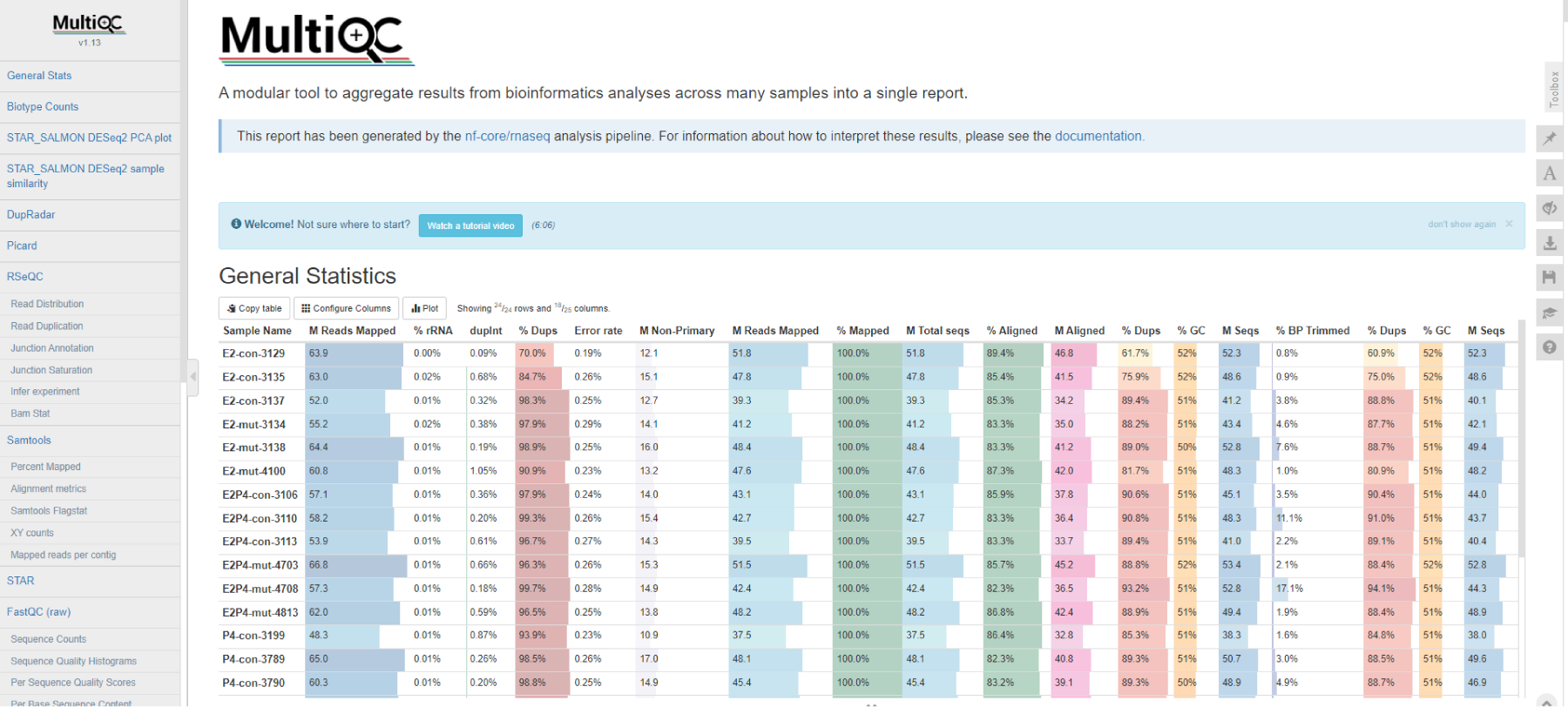
Screenshot of MultiQC report. The report summarizes quality control results of fastq files before and after each process.

### Midstream analysis with R

10. Read in rds object generated in the last step.

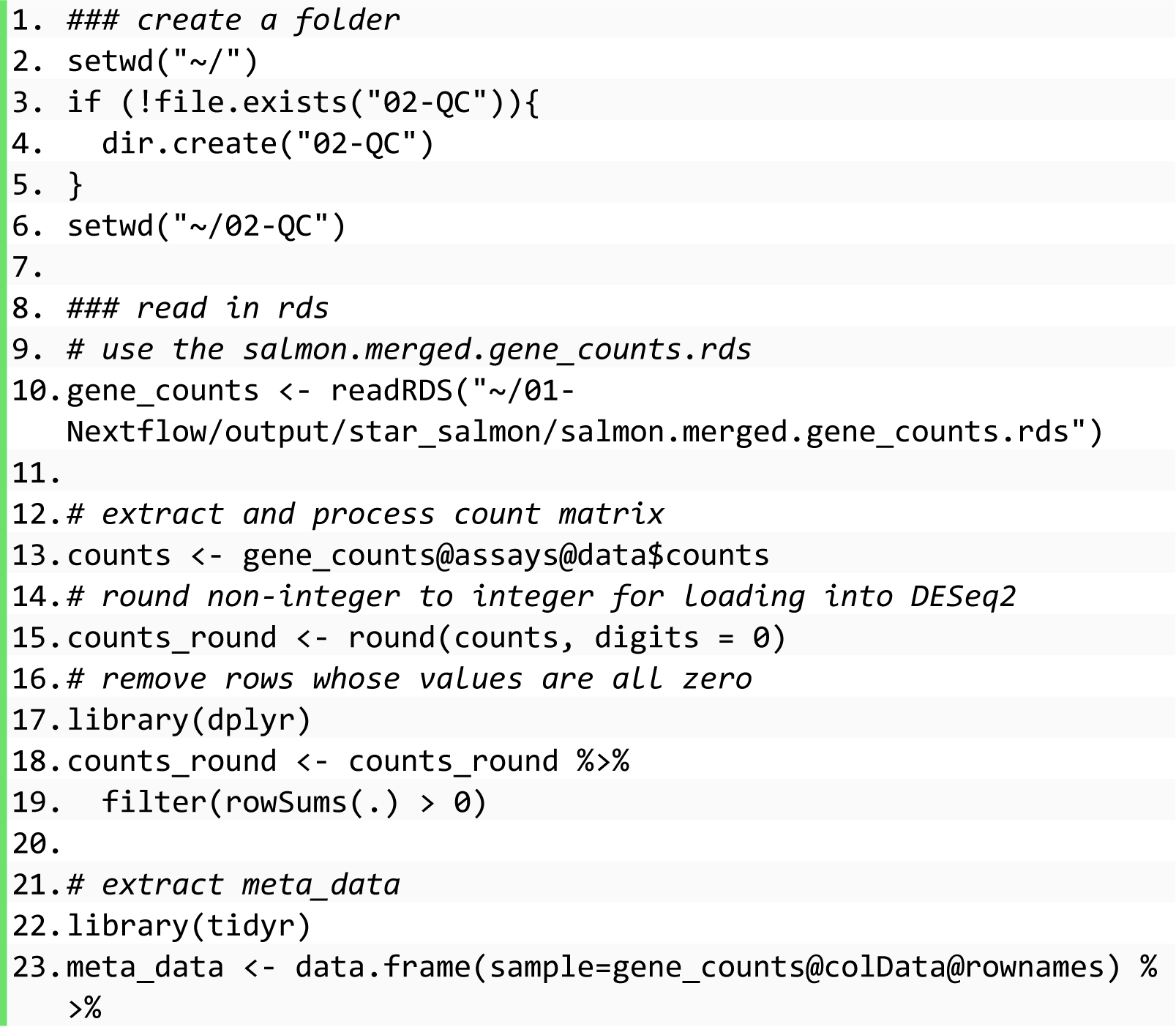

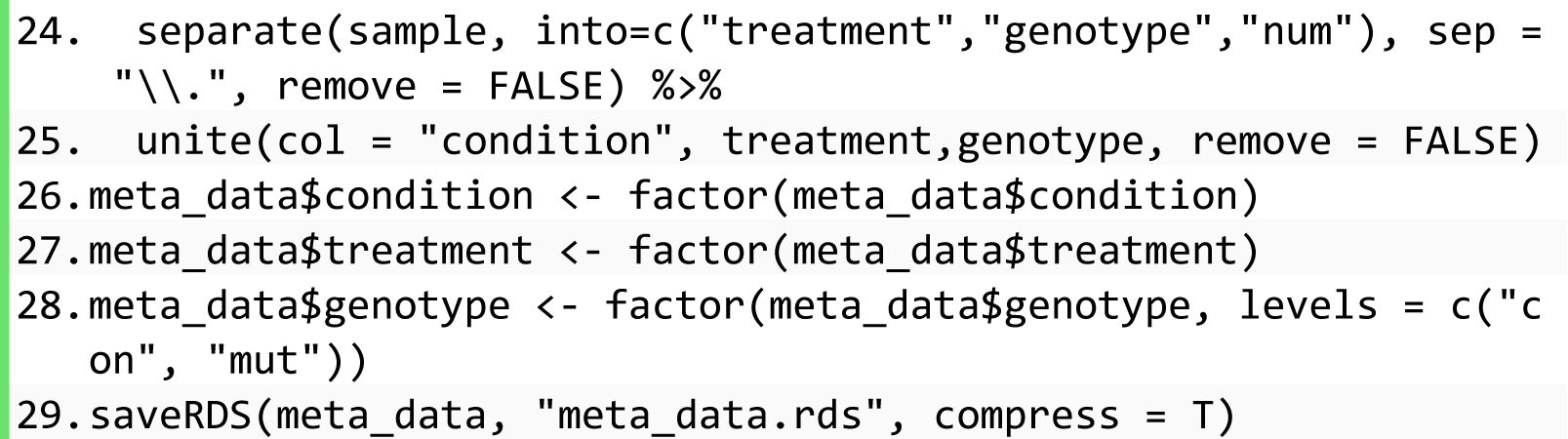
11. Generate a dds object with the DESeq2 R package.

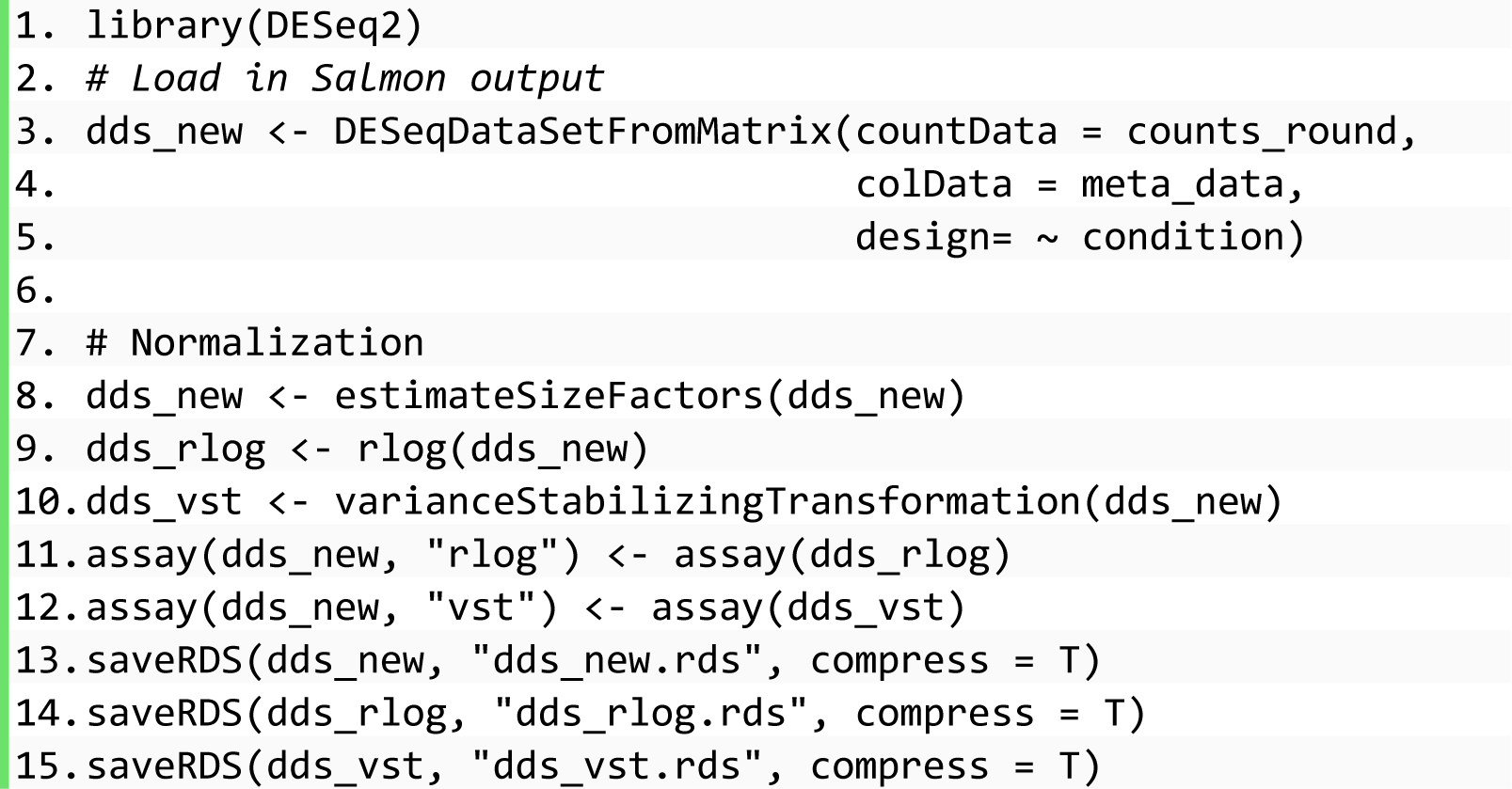
12. Plot to visualize the read counts.

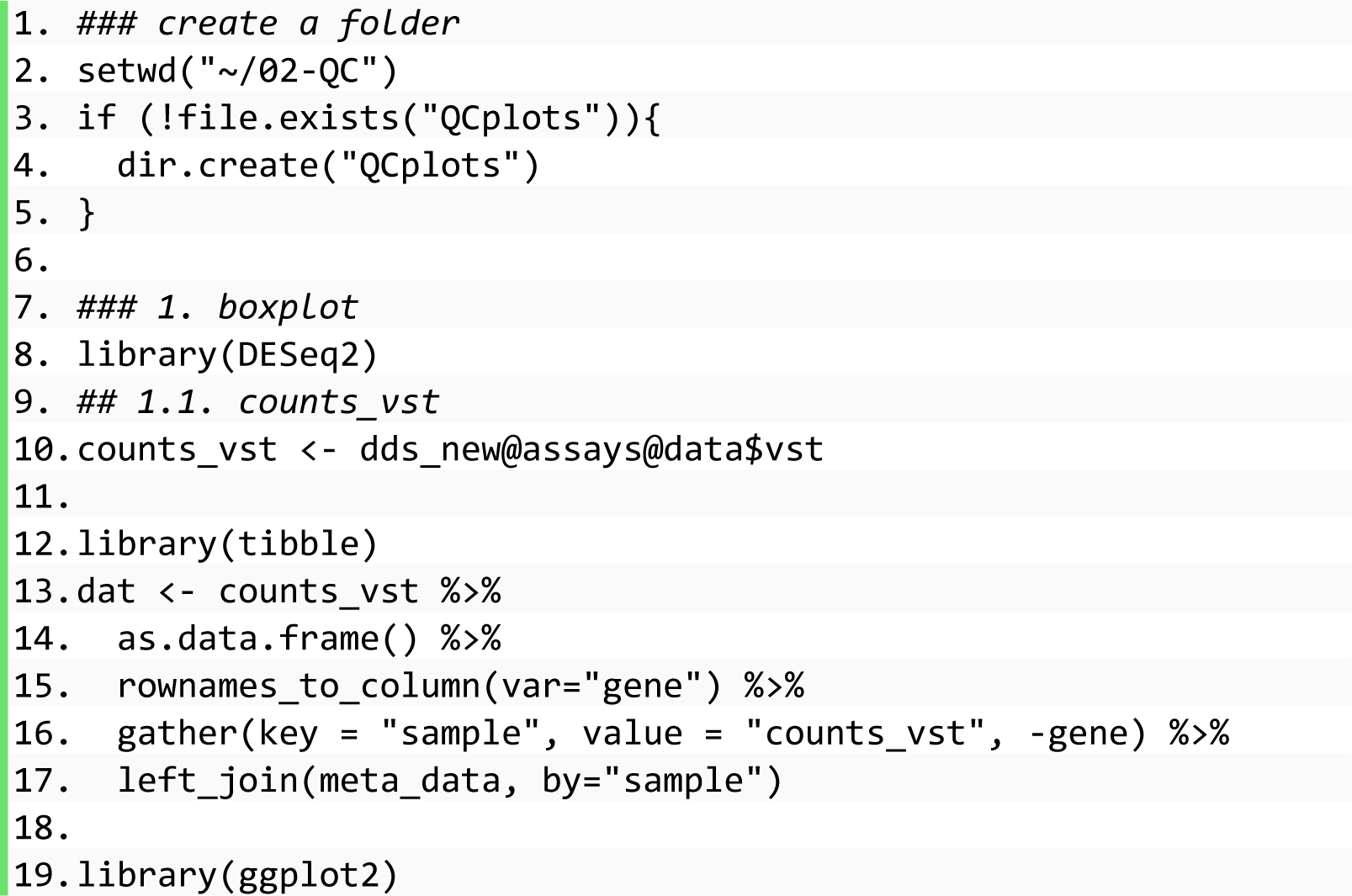

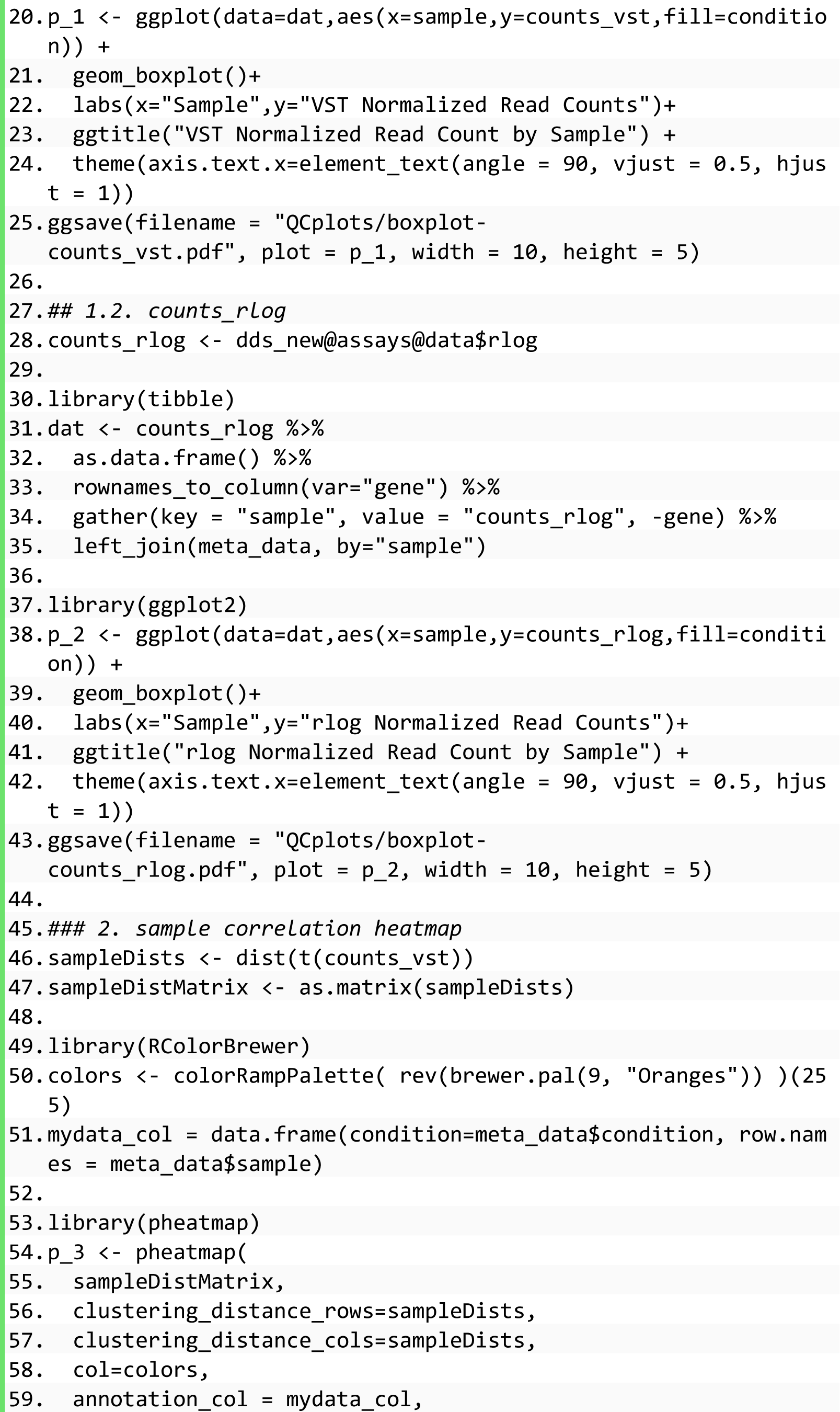

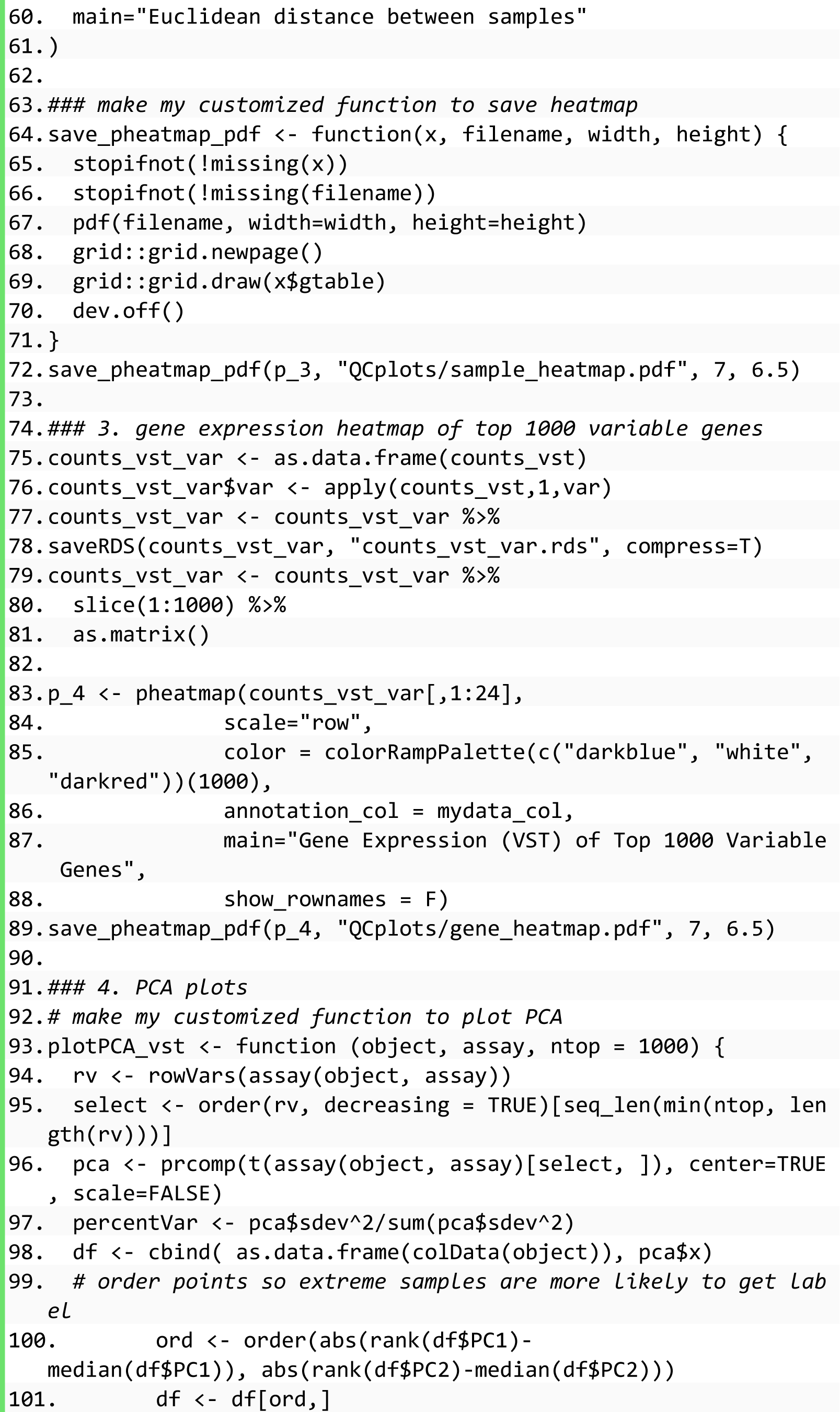

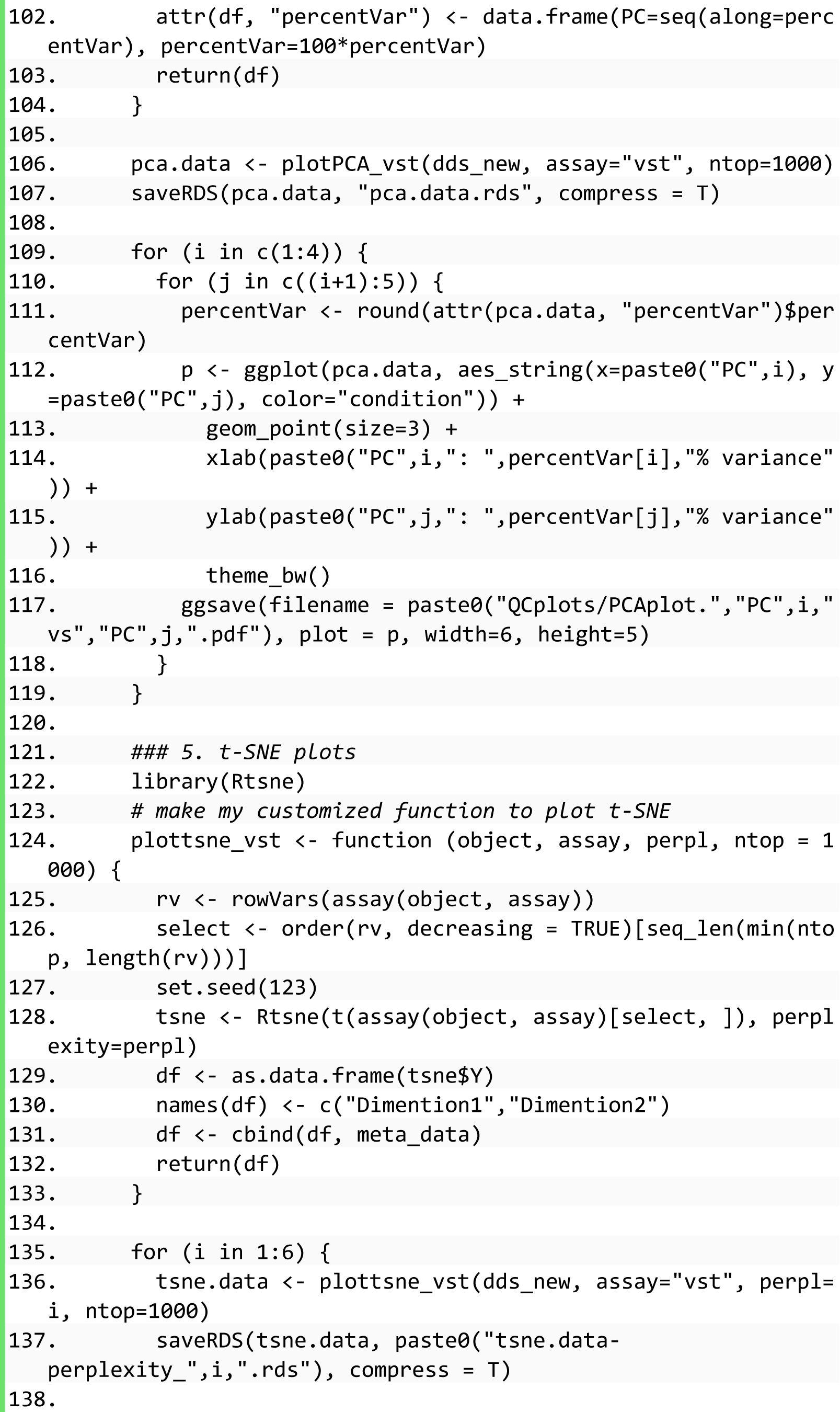

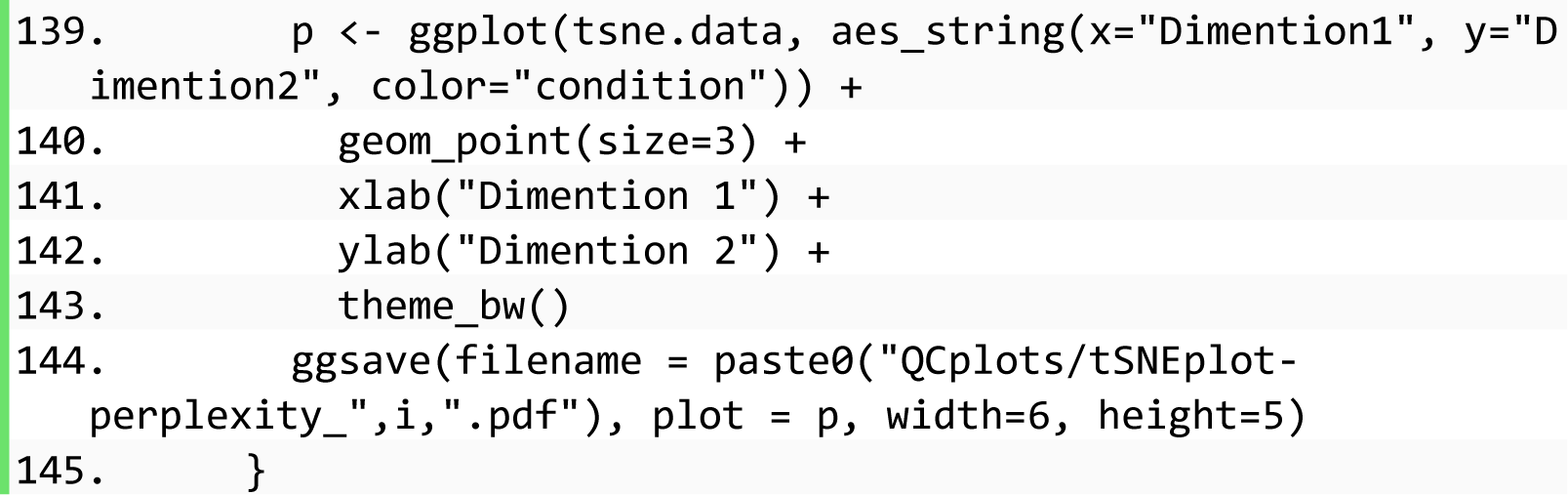
13. Visualization results of read counts. The raw data in the count matrix were normalized via two methods, rlog and vst, and were displayed in two box plots (Figure 3 and 4).

**Figure 3.**
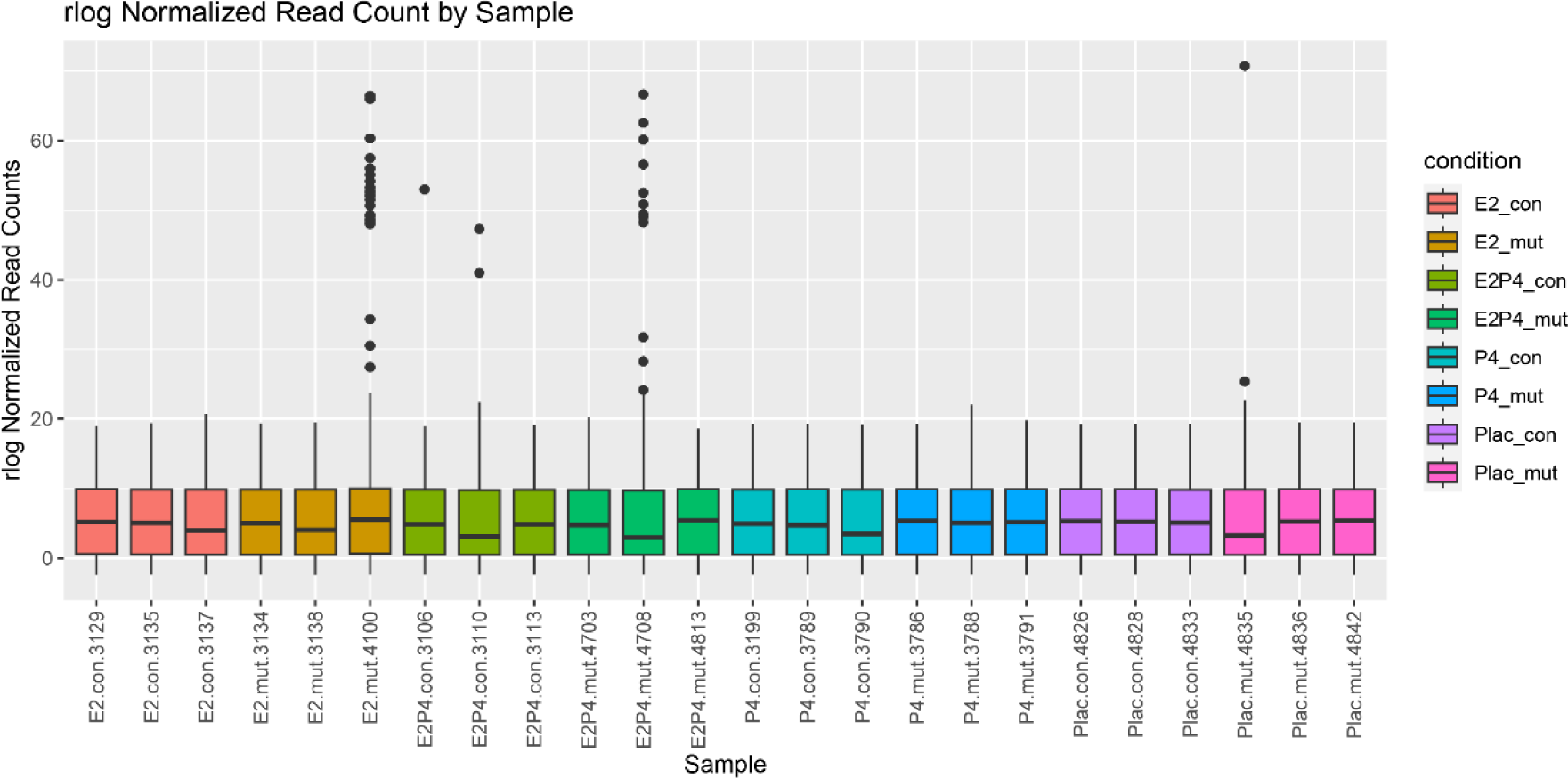
Box plot of read counts normalized by the rlog method. Samples in different groups are in different colors. The X-axis represents each sample. The Y-axis represents read counts normalized by the rlog method.

**Figure 4.**
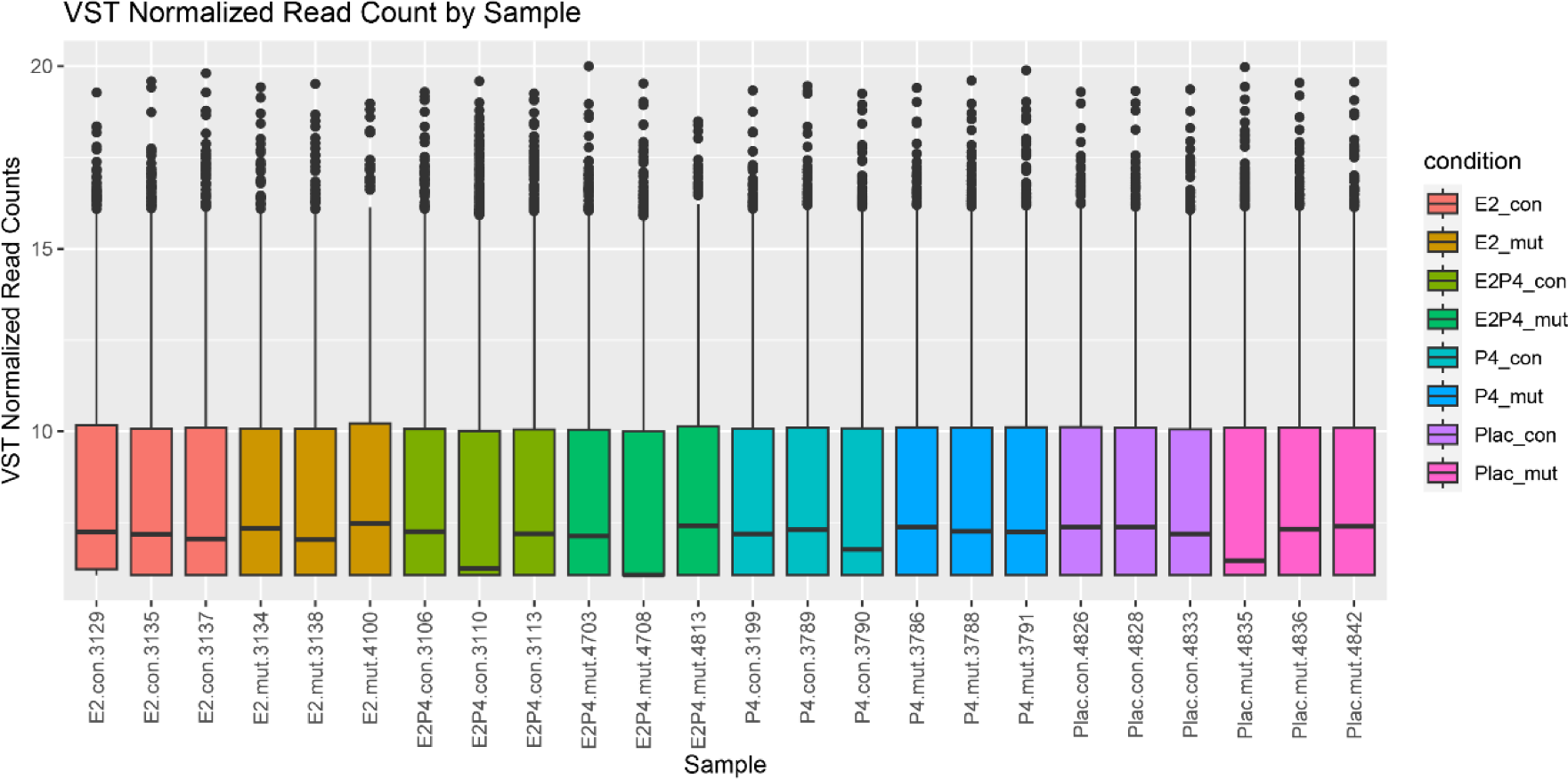
Box plot of read counts normalized by the vst method. Samples in different groups are in different colors. The X-axis represents each sample. The Y-axis represents read counts normalized by the vst method.

To observe the correlation among samples, the distance between each pair of samples was calculated and the Euclidean distance was visualized in the heatmap (Figure 5).

**Figure 5.**
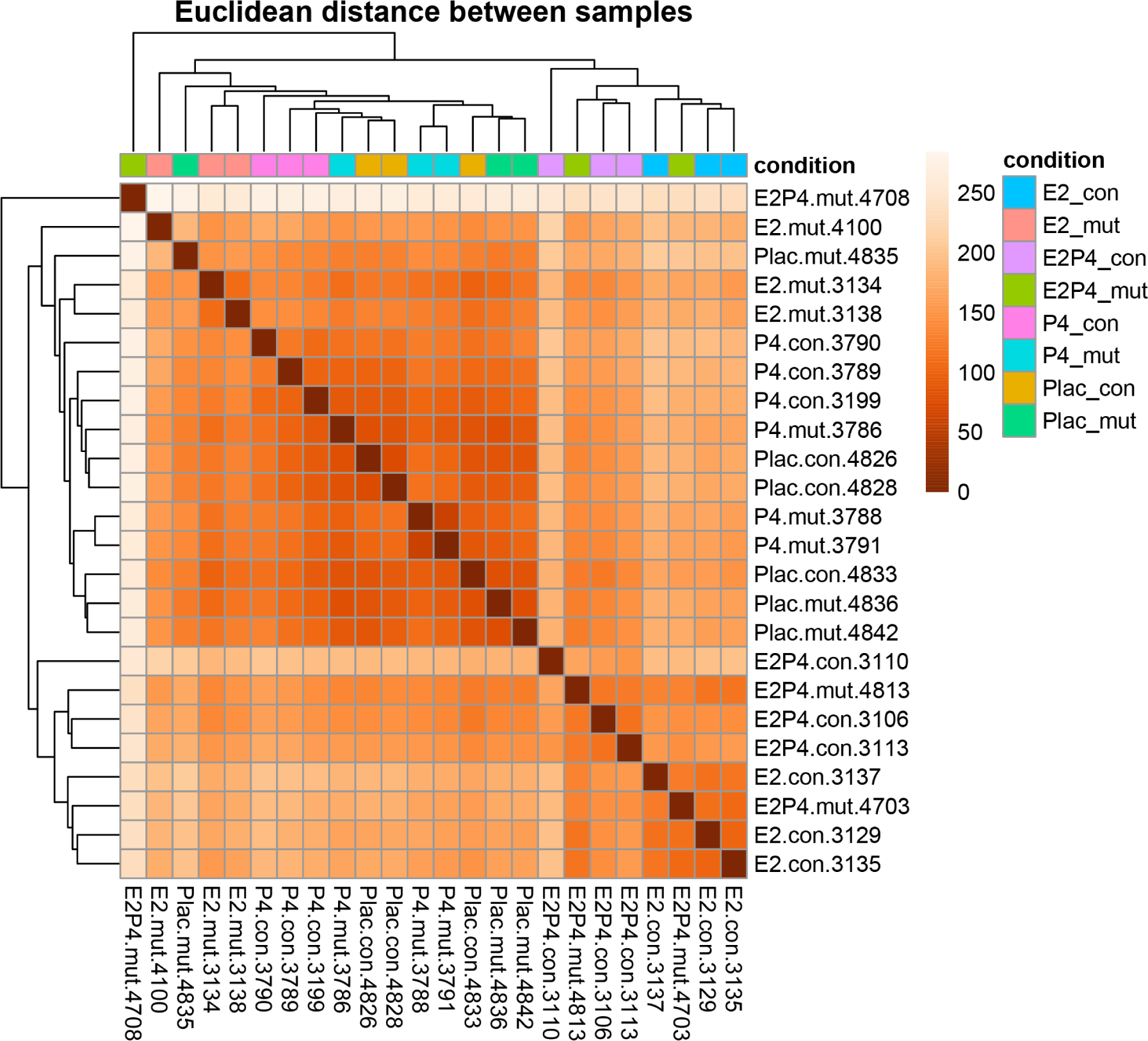
Heatmap of Euclidean distance between samples. Samples in different groups are in different colors. The color of the crossing square between two samples indicates the distance between those two samples. The darker the crossing square is, the smaller the distance between those two samples is.

In addition to the correlation at the sample level, the correlation across all samples was also calculated and visualized at the gene expression level (Figure 6).

**Figure 6.**
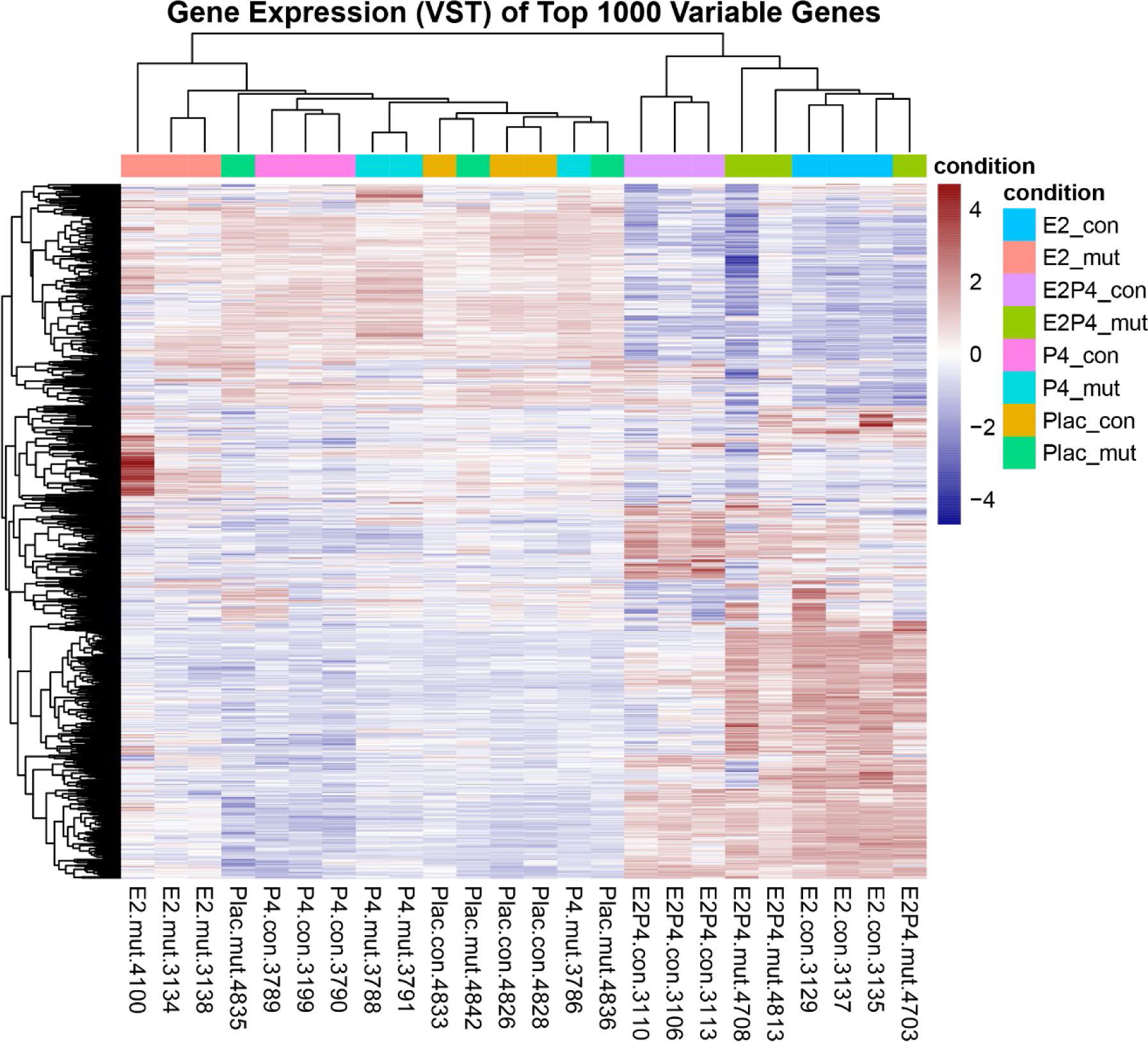
Heatmap of gene expression across samples. Samples in different groups are in different colors. The color of each gene indicates the scaled level of gene expression across samples. The redder indicates the higher level while the bluer indicates the lower level.

To further observe the distribution of samples, the dimension of read counts was reduced via principal component analysis (PCA) and visualized the results within combinations of each two of the first five principal components, i.e., PC1 and PC2, PC1 and PC3, PC1 and PC4, PC1 and PC5, PC2 and PC3, PC2 and PC4, PC2 and PC5, PC3 and PC4, PC3 and PC5, PC4 and PC5. The first two PCA plots were displayed here as examples (Figure 7A and 7B). In addition, the read counts were projected into the t-distributed Stochastic Neighbor Embedding (t-SNE) projection with a perplexity value from 1 to 6. The first and the last t-SNE plots were displayed here as examples (Figure 7C and 7C). Both PCA and t-SNE plots demonstrated the separation of samples by genotype and hormone treatment. Particularly, the E2 treatment affected gene expression the most significantly, and the P4 treatment regulated gene expression in a strikingly different way from the E2 treatment, while the inner group heterogeneity could not be ignored. The correlation observed in PCA and t-SNE plots was consistent with that in heatmaps at the sample distance level (Figure 5) and gene expression level (Figure 6).

**Figure 7.**
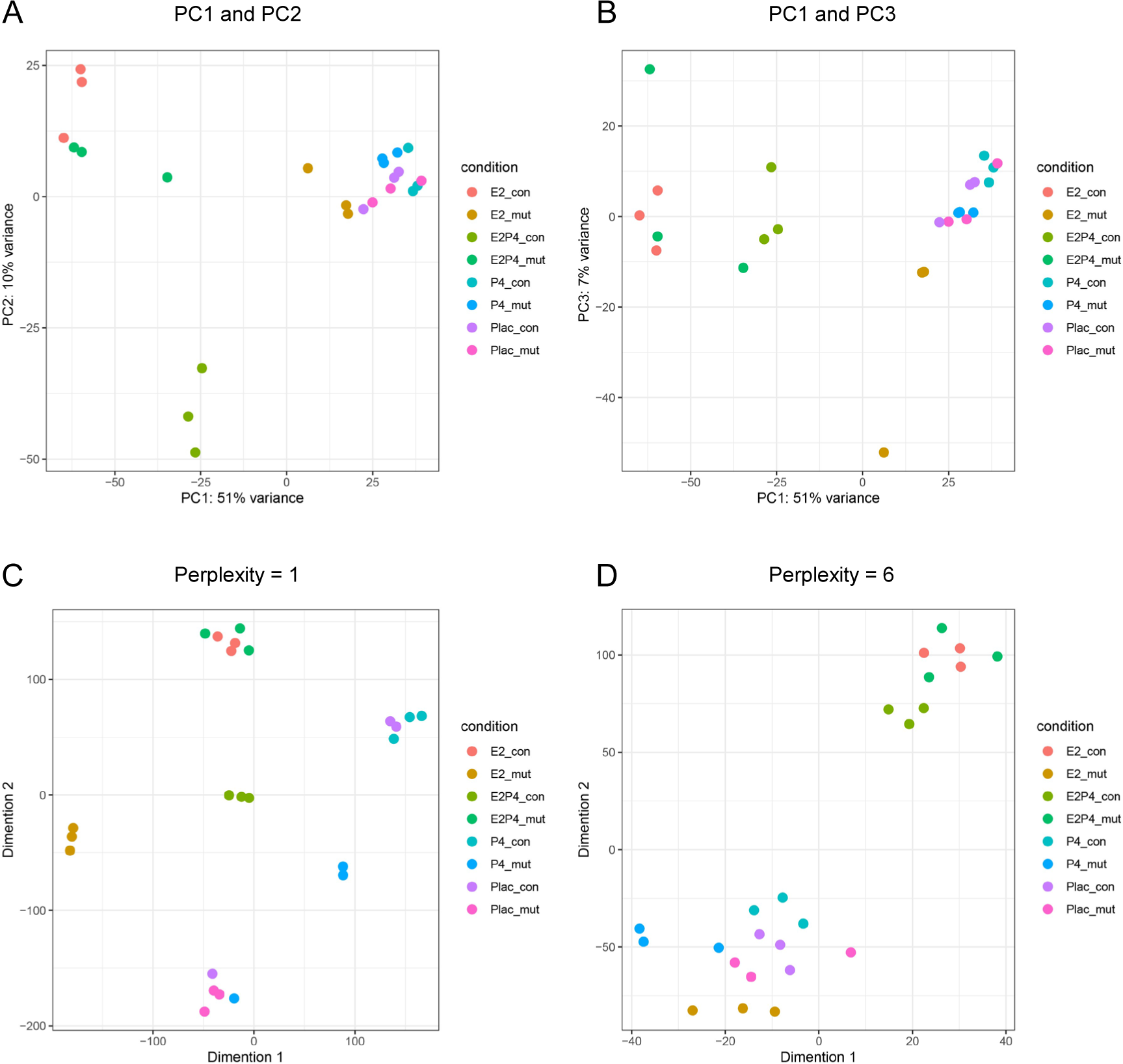
Sample distribution in projections of reduced dimension. A-B. PCA plots with PC1 and PC2 (A), and PC1 and PC3 (B). C-D. t-SNE plots with perplexity = 1 (C) and perplexity = 6 (D).

14. Convert gene ID from gene symbol to ENTREZID. The default gene IDs in the STAR output are gene symbols that are intuitive and convenient to read and understand. However, ENTREZID is preferred by many advanced bioinformatic software[18] because it is unique and more accurate than gene symbols. Therefore, I converted gene IDs from gene symbols to ENTREZIDs, which were required in subsequent analyses.

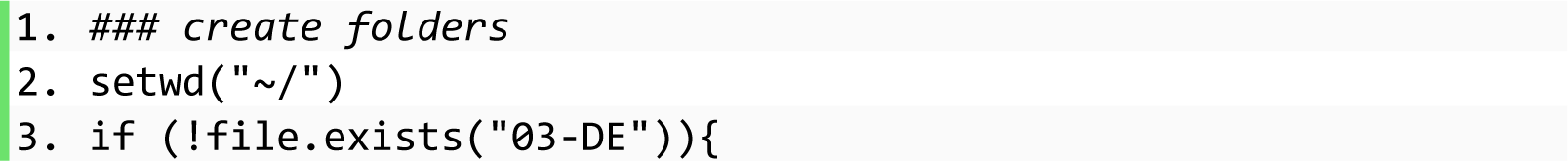

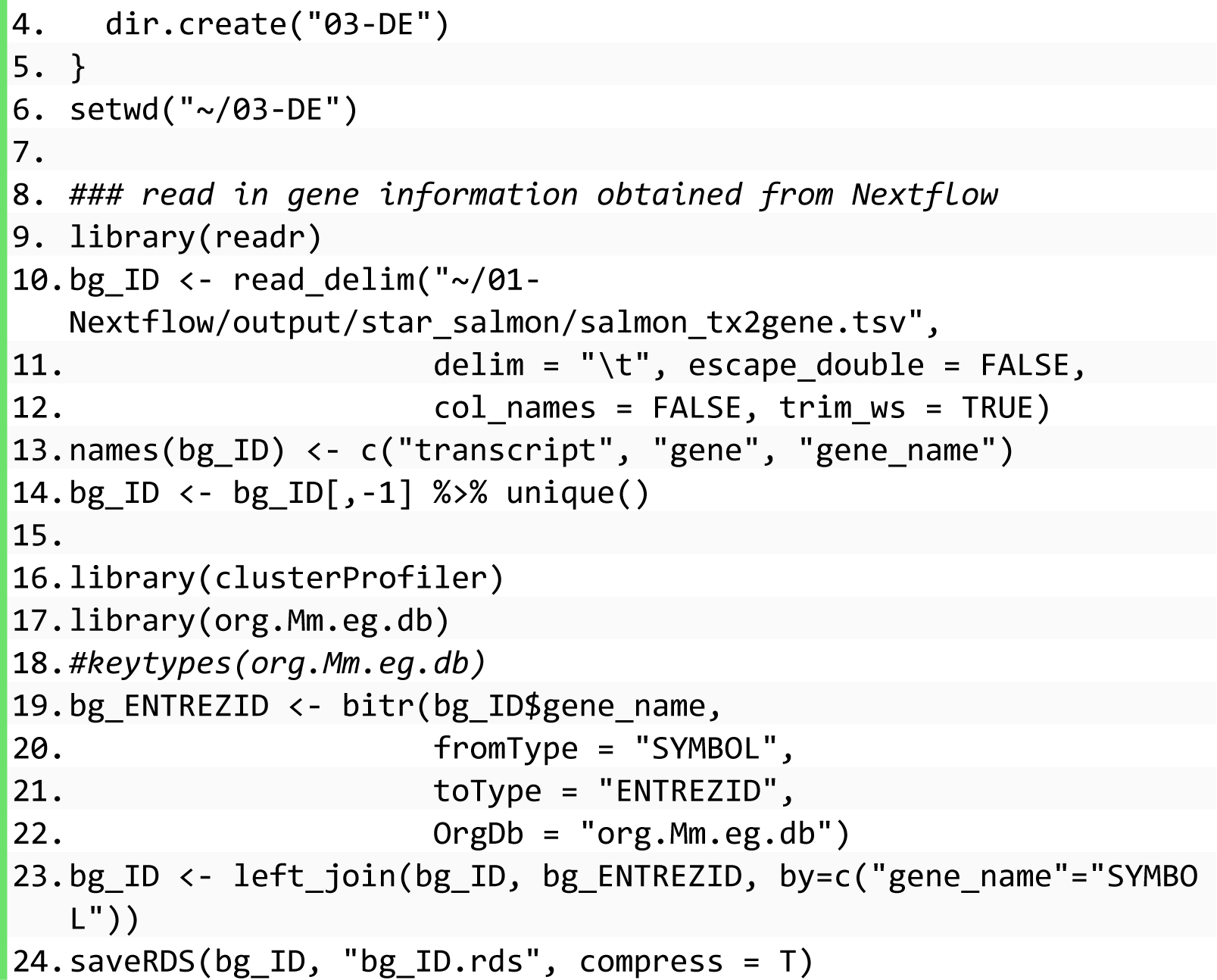
15. Differentially expressed gene (DEG) analysis. In this project, we aimed to compare the difference between mutant mice (mut) and their wild-type littermates (con) in each condition (hormone treatment or placebo) separately, but the same analysis codes can be used to compare every two groups of samples with minor adjustments.

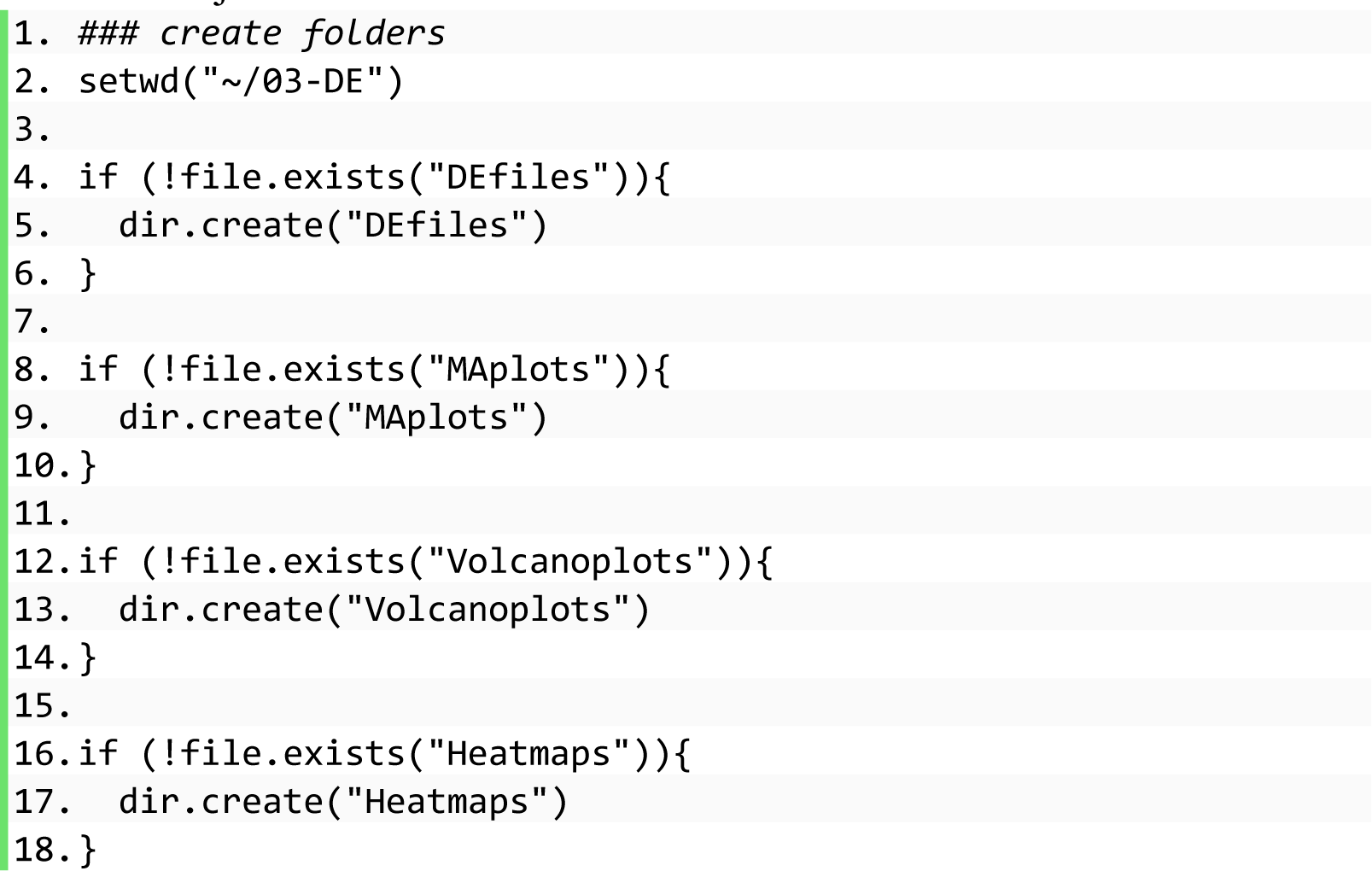

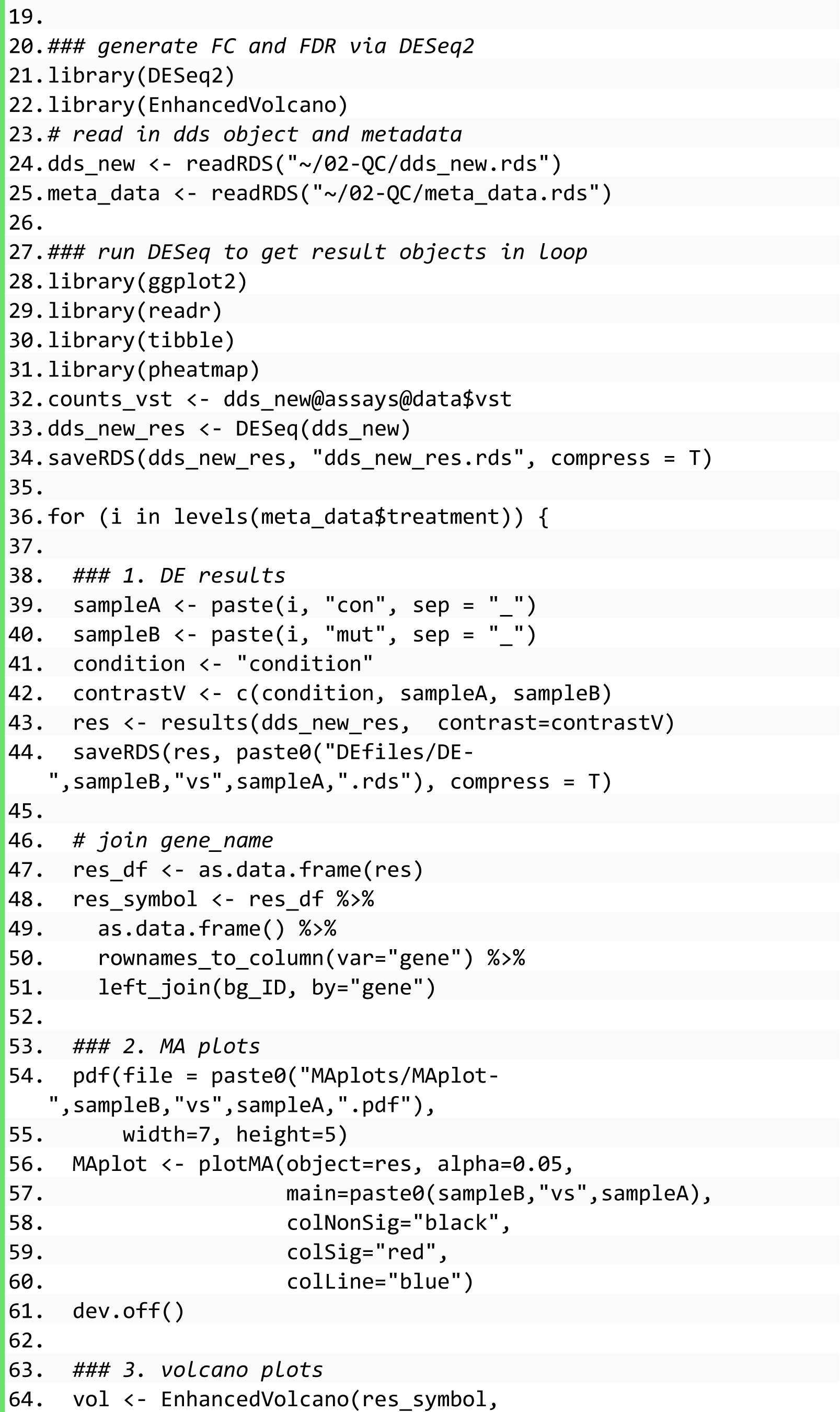

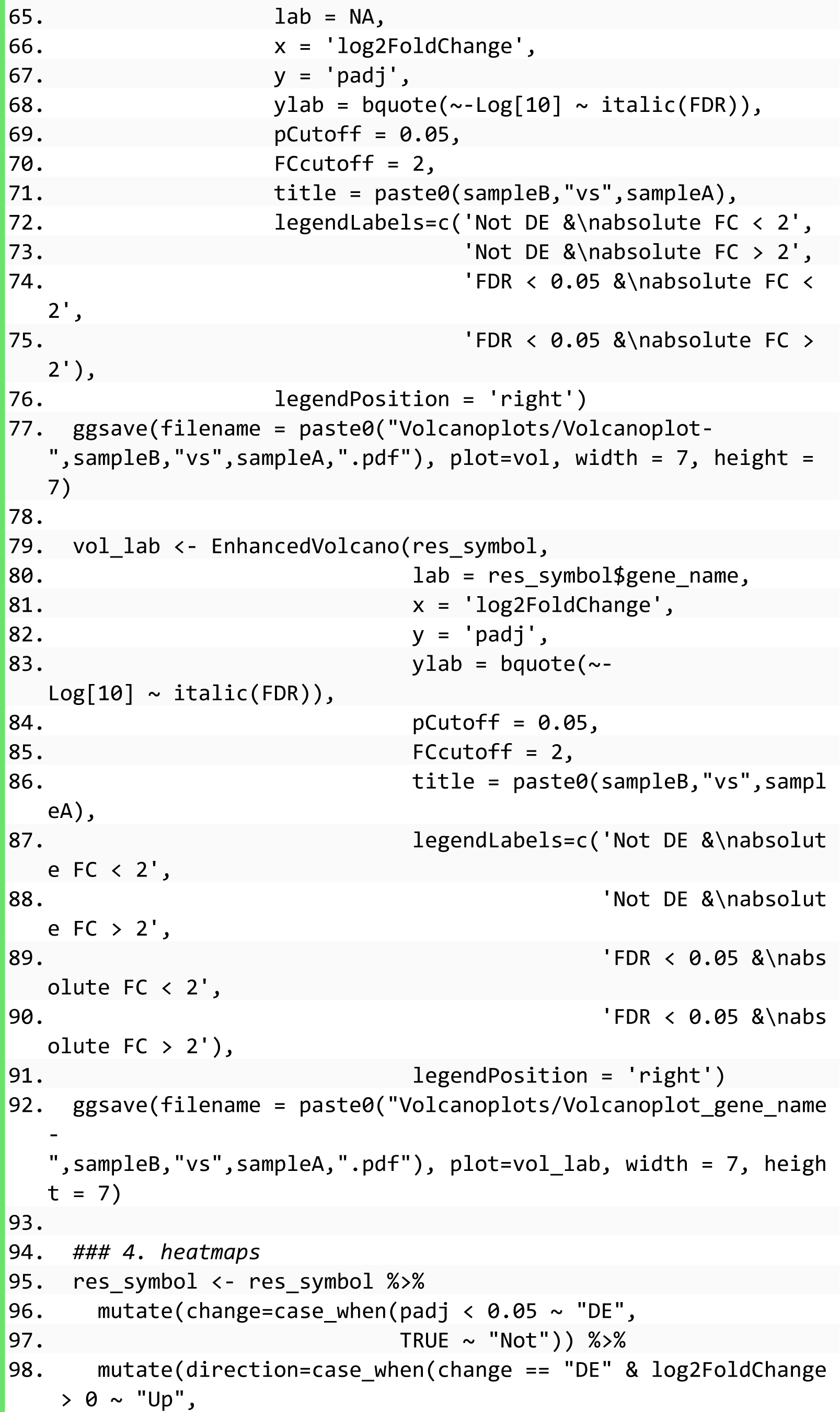

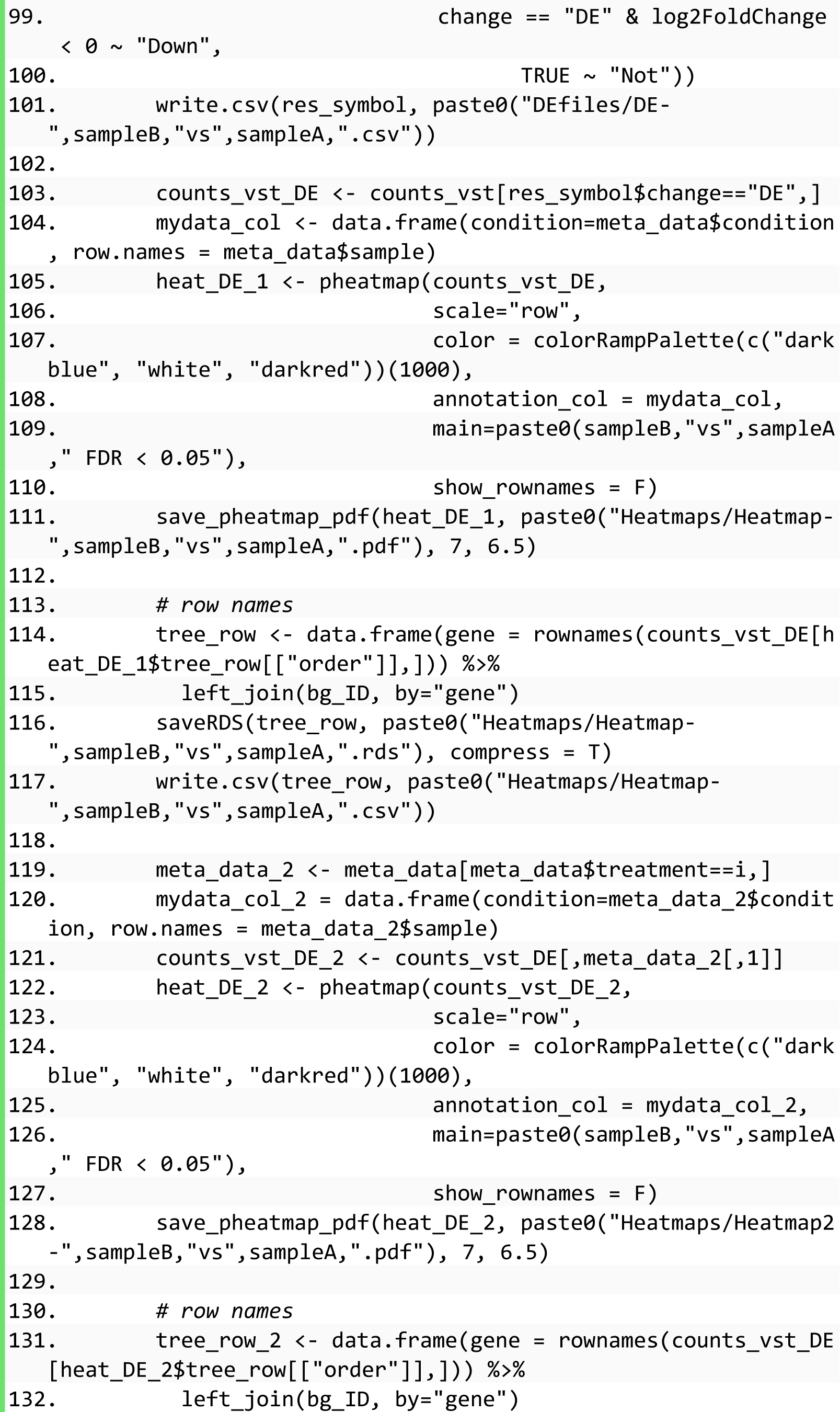

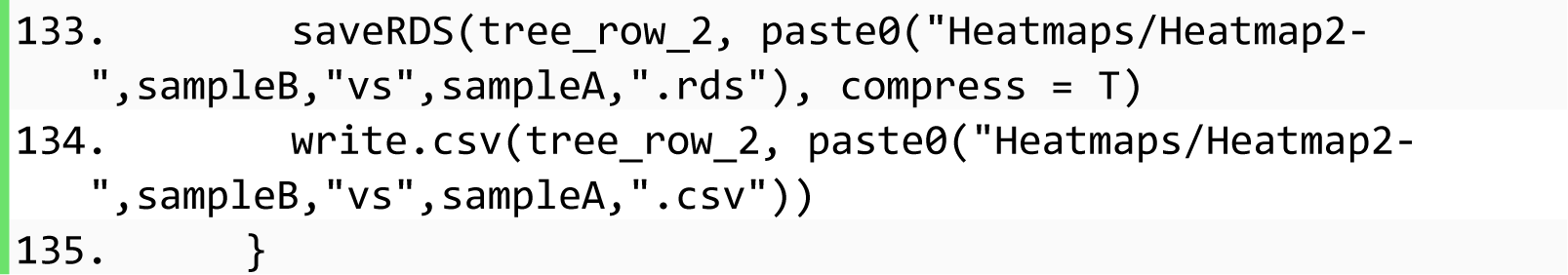
16. Visualization results of DEG analysis. Four sub-folders, i.e., Plac_mutvPlac_con, E2_mutvsE2_con, P4_mutvsP4_con, and E2P4_mutvsE2P4_con, were created inside the folder of ∼/03-DE/DEfiles containing results of DEG analysis. In addition, the results were visualized in various plots that were saved into sub-folders. MA plots suggested that the fold changes of genes between mut and con were distributed evenly and independent of mean values of normalized counts (Figure 8).

**Figure 8.**
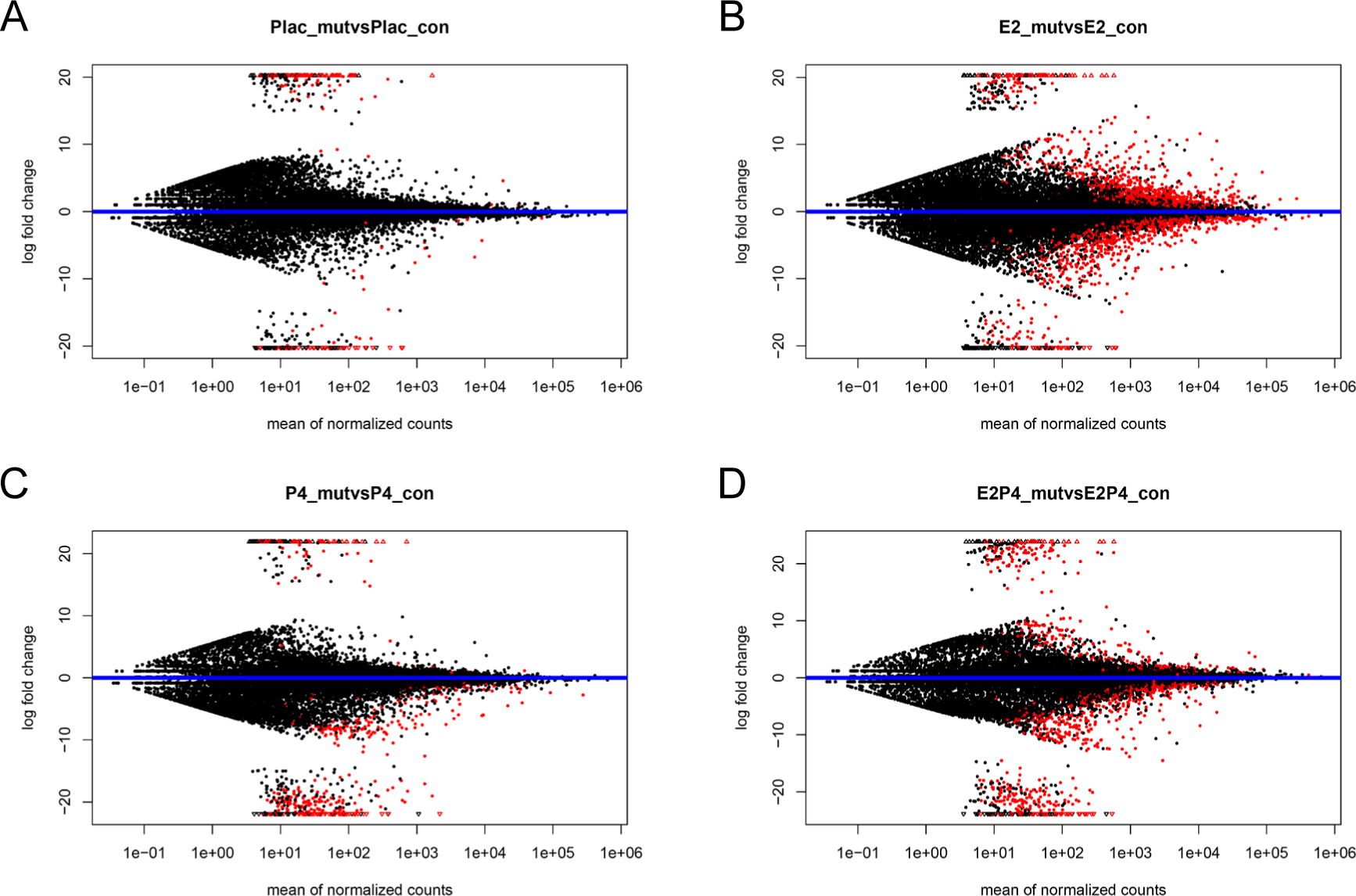
MA plots of DEG analysis results. X-axis represents the mean values of vst normalized counts. The Y-axis represents the log-transformed fold change in mut vs. con in each condition of hormone treatment. In addition to MA plots, volcano plots were also generated to observe the significant changes between mut and con. Because the above analyses suggested that E2 treatment had the most obvious effect on mut vs. con, the volcano plots of DEGs in mut vs. con under the E2 treatment were displayed here as examples (Figure 9). The panel on the right also labeled symbols of the most significantly changed genes (Figure 9B).

**Figure 9.**
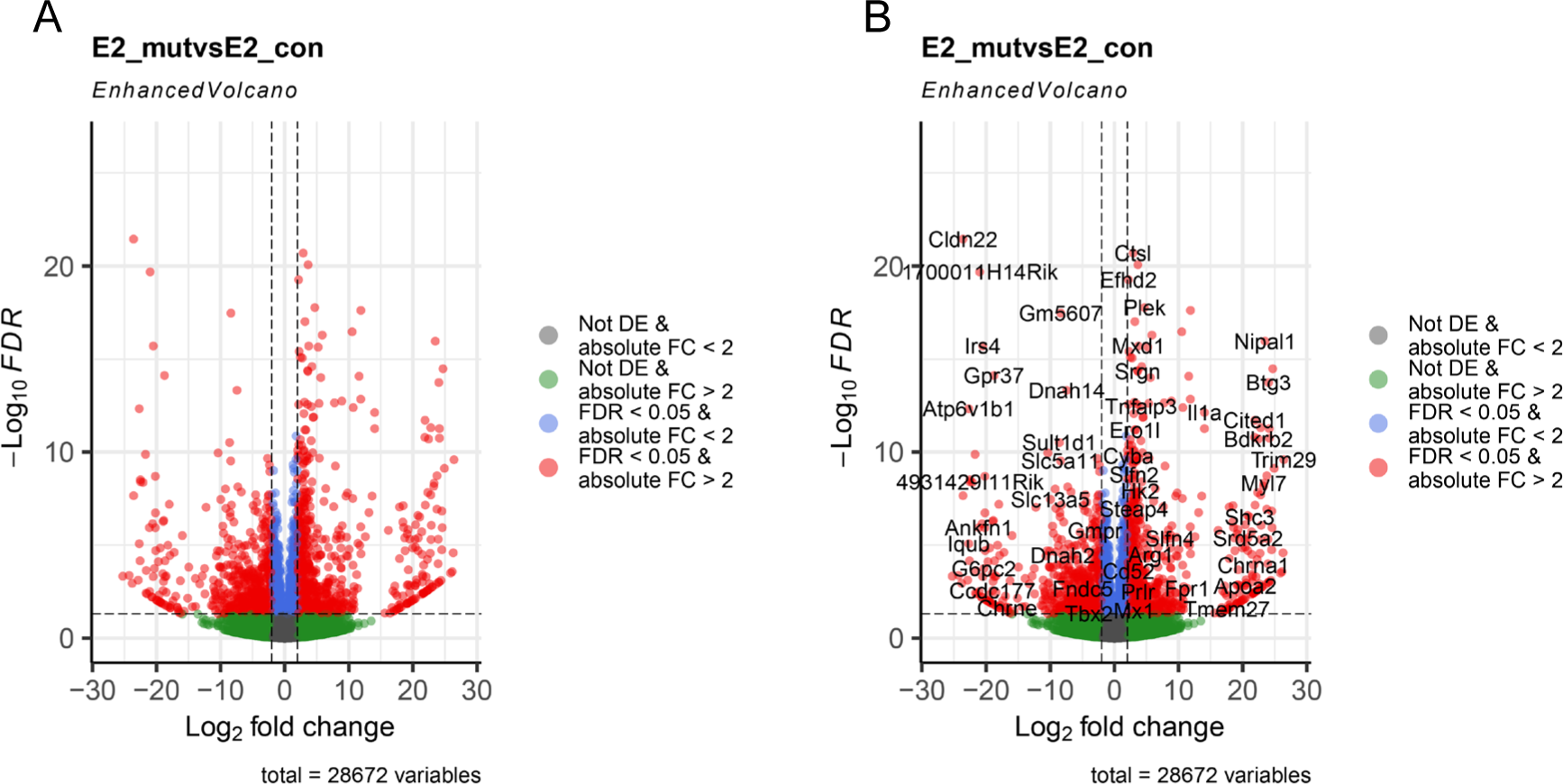
Volcano plots of DEGs in mut vs. con under the E2 treatment. The X-axis represents the log2 transformed fold change in mut vs. con. The Y-axis represents the minus log10 transformed FDR. Each dot represents one gene and is colored corresponding to the cutoffs shown in the legend. To observe the gene expression across samples, heatmaps were also generated for two groups of samples (Figure 10A) and all groups of samples (Figure 10B). The heatmap showed the two distinct clusters of genes that were upregulated or downregulated in E2_mut vs. E2_con (Figure 10A).

**Figure 10.**
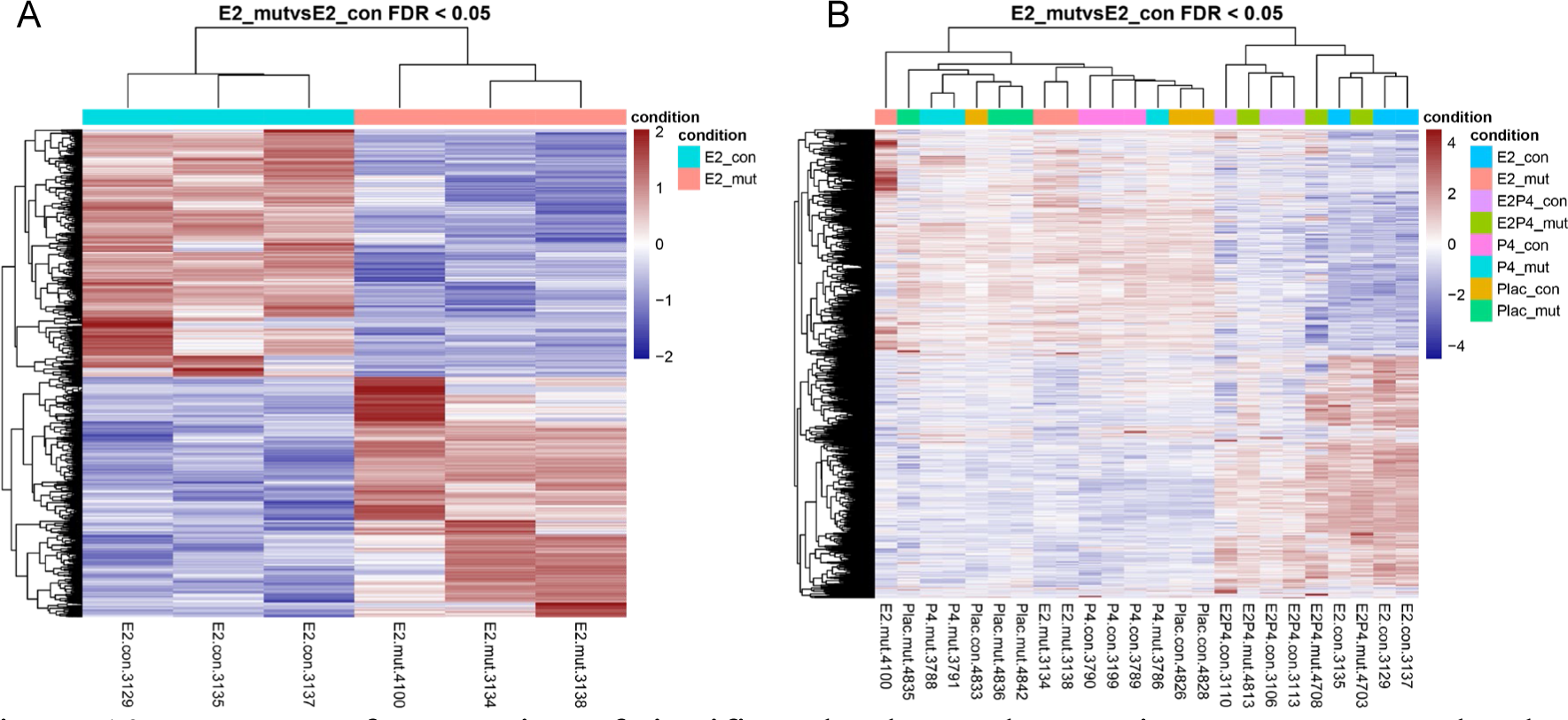
Heatmap of expression of significantly changed genes in mut vs. con under the E2 treatment. A. Heatmaps of comparison between two groups of samples. B. Heatmap across all samples. Samples in different groups are in different colors. The color of each gene indicates the scaled level of gene expression across samples. The redder indicates the higher level while the bluer indicates the lower level.

### Downstream analysis with R

17. Functional analyses including Gene Ontology (GO) – Biological Process (BP), Cellular Component (CC), and Molecular Function (MF), and KEGG analyses.

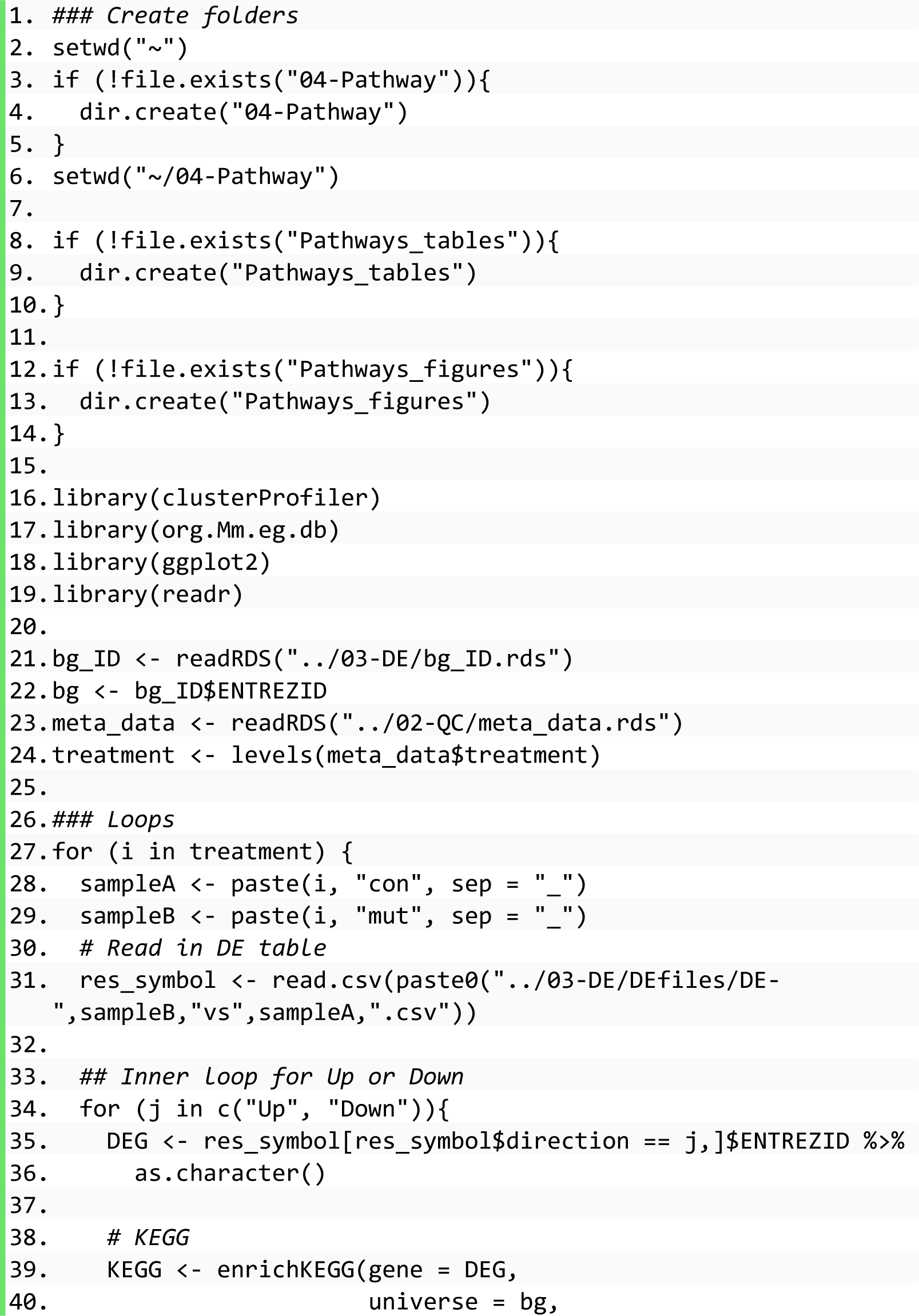

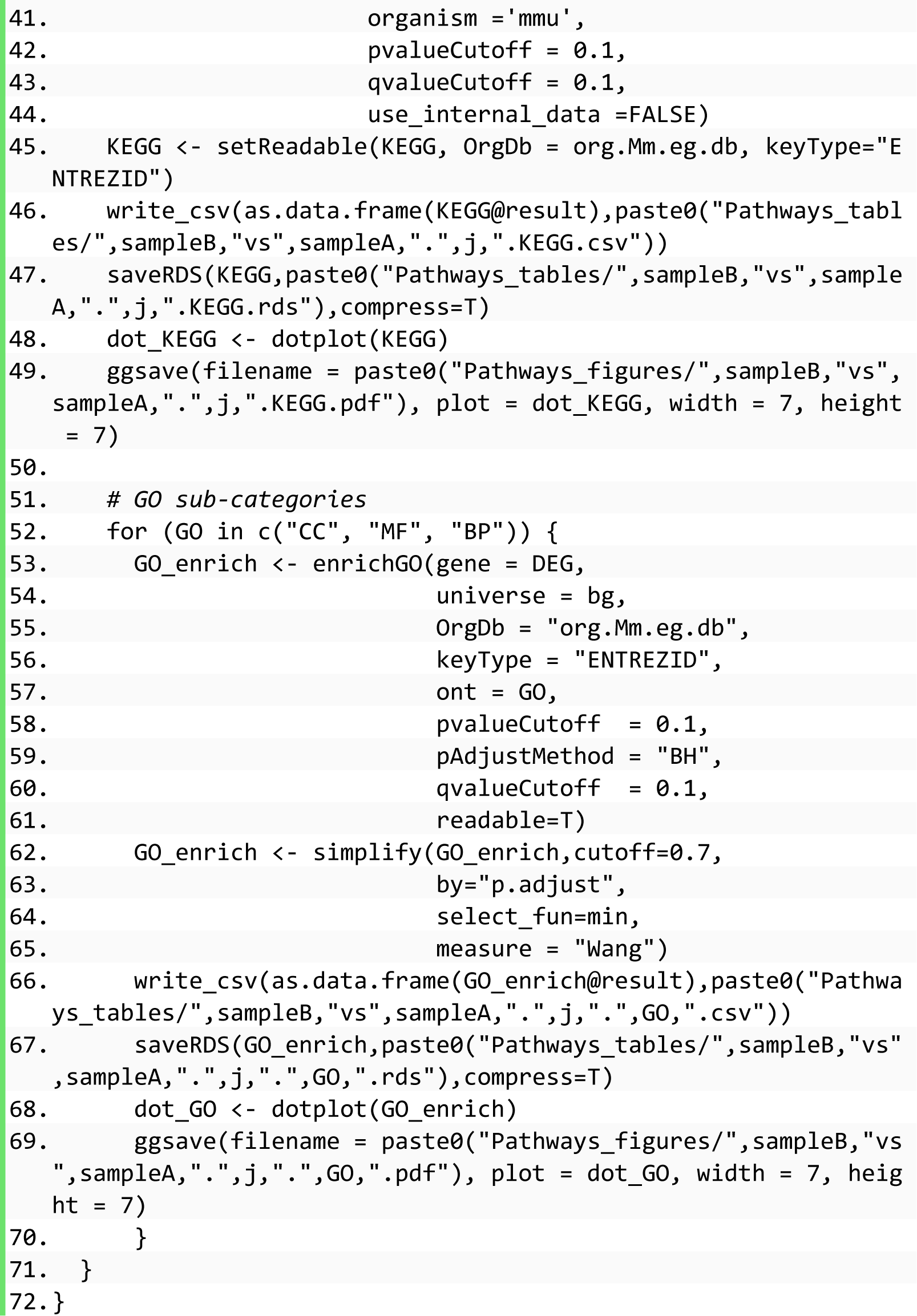
18. Visualization of results of functional enrichment analysis. DEGs were divided into two parts, i.e., upregulated (Up) and downregulated (Down) genes in mut vs. con under individual hormone treatment. The two parts were separately loaded into the clusterProfiler R package for functional enrichment of KEGG pathways and GO terms including BP, MF, and CC sub-categories. Here, the bubble charts of KEEG pathways and GO terms enriched from DEGs in mut vs. con under the E2 treatment were displayed as examples (Figure 11).

**Figure 11.**
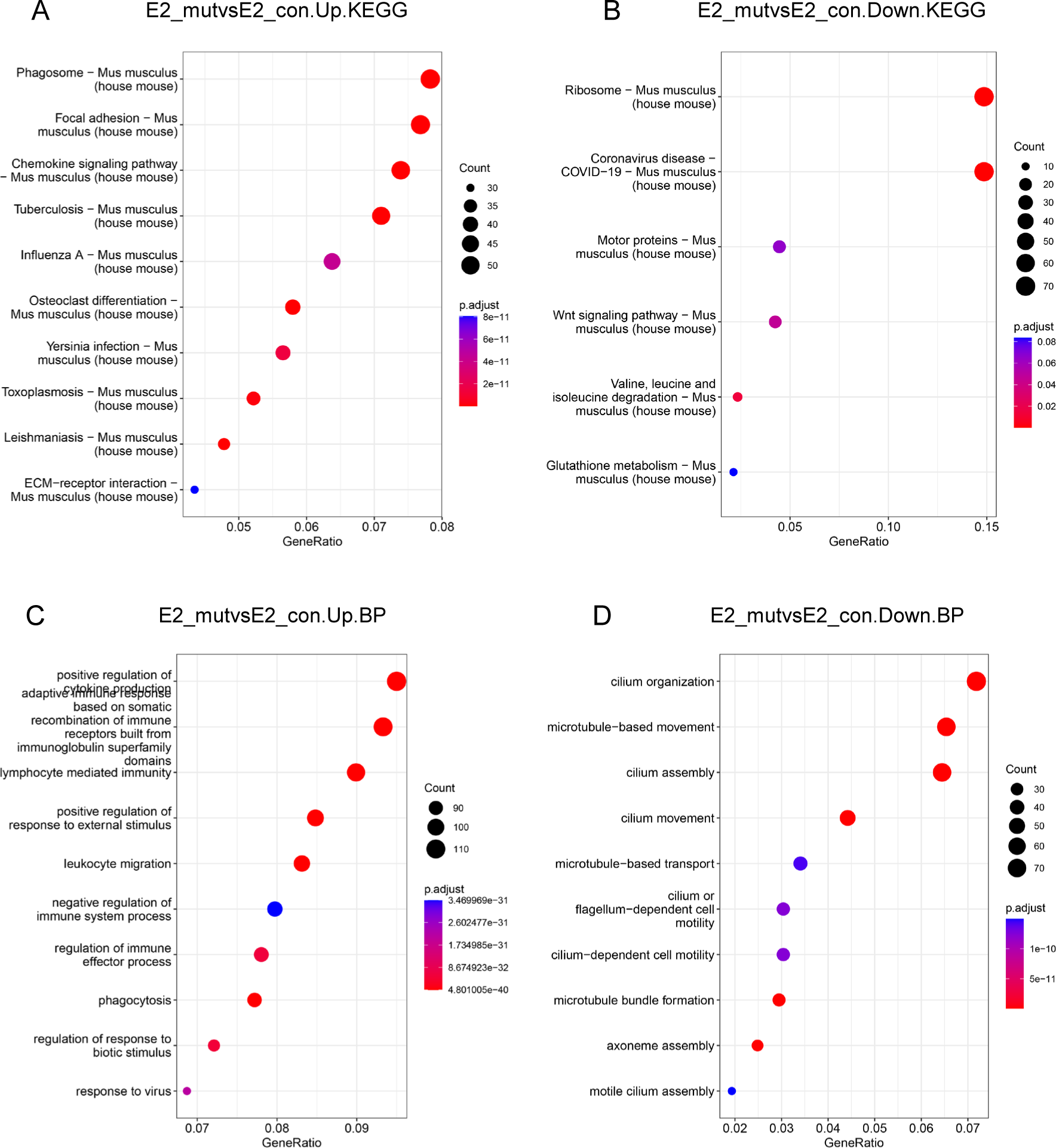
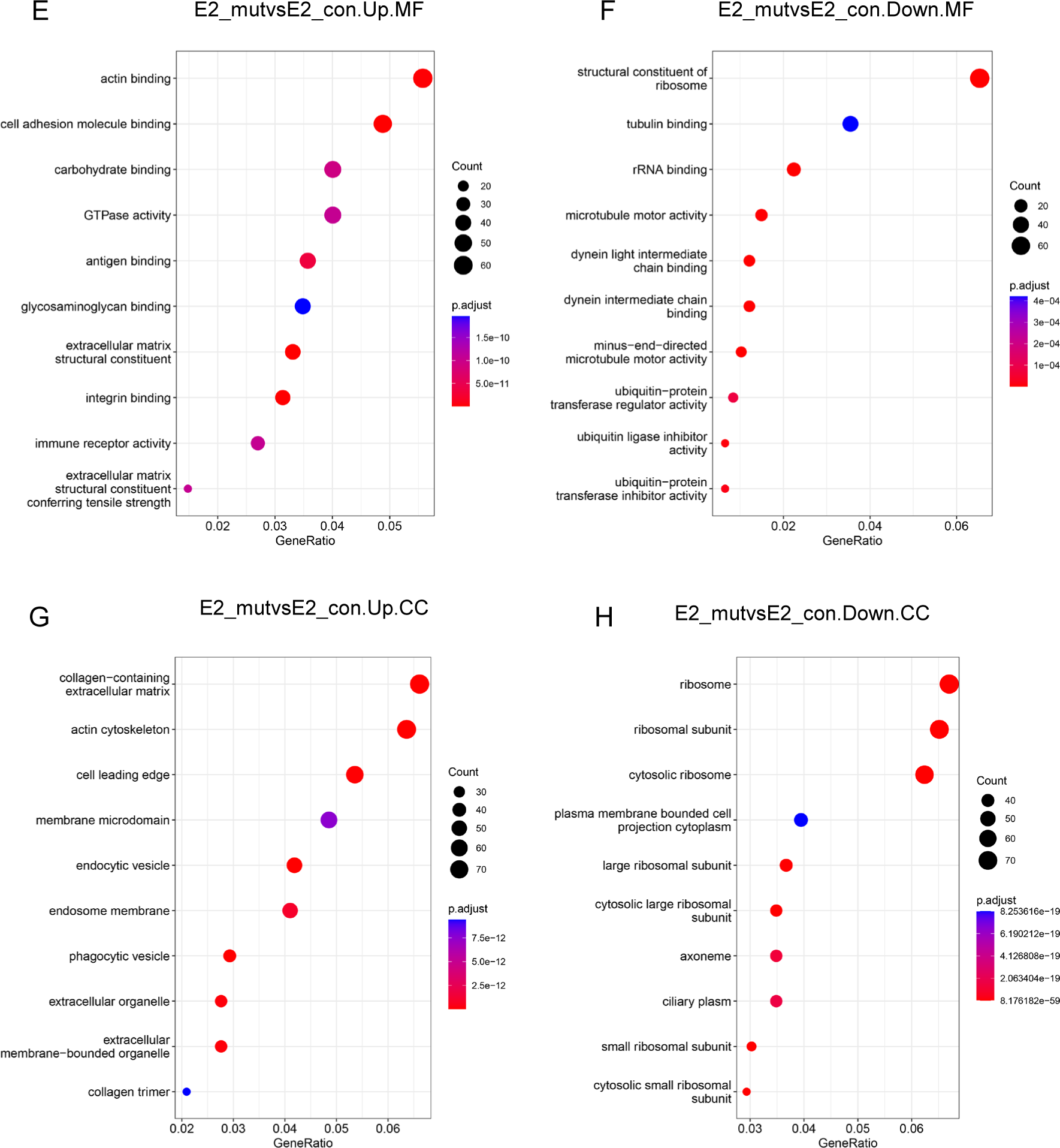
Functional enrichments. IDs of significantly upregulated and downregulated genes in E2_mut vs. E2_con were converted into the ENTREZ IDs and imported into the clusterProfiler R package to conduct functional enrichment. The top ten terms with adjusted p-values < 0.05 were displayed in bubble charts. Four categories were KEGG pathways from the KEGG pathway database (A and B), GO_BP terms from the Biological Process (Gene Ontology) database (C and D), GO_MF terms from Molecular Function (Gene Ontology) database (E and F), and GO_CC from Cellular Component (Gene Ontology) database (G and H). Color represents significance: the redder, the more significant.

## Discussion

This pipeline started from processing raw fastq files obtained from the Illumina HiSeq4000 sequencing machine, aligned reads to the Ensembl/GRCm38 reference genome via STAR, quantified to count matrix via Salmon, visualized normalized counts, analyzed and visualized DEGs, enriched and visualized KEGG pathways and GO terms to deduce changed functions in R.

This pipeline combined the advantages of Nextflow and R from upstream to midstream, and to downstream analyses. In the upstream analysis, different reference genomes can be downloaded from the igenomes repository (https://support.illumina.com/sequencing/sequencing_software/igenome.html) provided by Illumina. In addition to STAR and Salmon, other alignment and quantification software can be selected as a substitution. Addition parameters can be adjusted in the Nextflow config file. Therefore, this pipeline that was built based on Nextflow can be extended to analyze RNA-seq data generated from other species and with different qualities.

In the midstream and downstream analyses, this pipeline took advantage of R in statistics and visualization. In addition to the pairs of comparisons that were demonstrated here as examples, samples from any two groups can be compared via DESeq2 by this pipeline. I am considering expanding this pipeline to provide other options such as EdgeR and Limma which are also widely used. Of note, this pipeline inherited one of the best advantages of R, i.e., the customized and fast visualization. I provided codes to observe the data and results in many most popular plots. The generated figures can be combined and trimmed flexibly. All figures can be exported to PDF files which are editable and can be transformed into any other formats in graph edition software like Illustrator.

In summary, this pipeline provides an easy and convenient way to analyze RNA-seq data and reproduce bioinformatics analysis. Particularly, the enriched KEGG pathways and GO terms are very useful in the interpretation of biological functions and the discovery of regulatory mechanisms. The results demonstrated that age-associated alterations of mutation and hormones reprogram the transcriptome significantly. In addition to the age-related factors studied in this project, this pipeline can also be broadly applied in other RNA-seq data analyses in the future.

## Notes

### Competing Interest Statement

The authors have declared no competing interest.

### Summary of Updates

Add affiliation

